# Methylome profiling of SetDB1 deficient ESCs reveals diverse epigenetic cross-talk during pluripotency

**DOI:** 10.1101/2025.06.18.660343

**Authors:** Nick G.P. Bovee, Stefan H.A. Hoogland, Ehsan Habibi, Siebren Frölich, Arie B. Brinkman, Klaas Mulder, Joop H. Jansen, Matthew Lorincz, Henk G. Stunnenberg, Hendrik Marks

## Abstract

SetDB1 is best known as a chromatin modifier catalyzing H3K9me3. However, recent studies show that SetDB1 can promote H3K27me3-deposition and CTCF-binding, and potentially recruit de novo DNA methyltransferases. Given the tight connection with these processes, we hypothesized that DNA methylation (DNAme) may integrate these combined features of SetDB1. Thereto, we conducted time-course whole-genome bisulfite sequencing following *Setdb1* knockout (KO) in mouse embryonic stem cells (ESCs). In serum-cultured ESCs, nearly half of SetDB1 binding sites are DNA methylated, coinciding with H3K9me3, mainly silencing retrotransposons and imprinting control regions. Both H3K9me3 and DNAme are lost upon *Setdb1* KO, but while TET2 rapidly removes DNAme at many of these sites, some retrotransposons are shielded from TET2 and lose DNAme slowly via passive dilution. SetDB1-mediated regulation via H3K27me3, CTCF, SMAD3, and histone acetylation are uncoupled from the DNAme-H3K9me3 axis. Hypomethylated naïve ESCs show massive reactivation of retrotransposons upon *Setdb1* KO, providing functional evidence that DNAme adds a protective layer against such activity. Altogether, our findings reveal how DNAme coordinates the multifaceted regulatory roles of SetDB1.

**Highlights:**

1. In serum ESCs, SetDB1-dependent deposition of H3K9me3 and DNAme are tightly coupled, primarily silencing repeats and imprinted control regions;
2. Loss of SetDB1 causes demethylation of a large range of repeat types, the pace of which is dependent on TET pre-loading;
3. The regulatory modes of SetDB1 mediated by H3K27me3, CTCF, TGF-β signalling and histone acetylation are uncoupled from the SetDB1 DNAme-H3K9me3 axis and/or from each other;
4. Loss of SetDB1-dependent DNAme in hypomethylated 2i ESCs reveals that DNAme serves as a buffering layer to repress SetDB1-mediated H3K9me3 targets.

## Introduction

Chromatin regulation plays an essential role throughout mammalian development, controlling gene expression to ensure proper cell fates. Extensive chromatin remodeling governs developmental processes, maintaining cell identity and function. In pluripotent mouse embryonic stem cells (ESCs), SetDB1, which targets up to twenty thousand genomic loci (Bilodeau et al., 2009; Karimi et al., 2011; Yuan et al., 2009), plays an essential role in regulating various cellular processes, including maintenance of the embryonic stem cell state, differentiation and metabolism (Bilodeau et al., 2009). Indeed, depletion of SetDB1 in mice is lethal during implantation (Dodge et al., 2004). SetDB1 is primarily recognized for its histone methyltransferase activity, which is mediated by its bifurcated SET domain that catalyzes trimethylation of lysine 9 on histone 3 (H3K9me3) (Markouli et al., 2021), a hallmark of heterochromatin. However, despite widespread deposition of this mark, depletion of SetDB1 leads to de-repression of only 33 H3K9me3-associated genes in ESCs (Karimi et al., 2011). This indicates that SetDB1 mainly facilitates H3K9me3-mediated silencing at non-genic regions (Matsui et al., 2010; Leung & Lorincz, 2012; Karimi et al., 2011).

Further complicating the characterization of SetDB1 function, nearly half (46%) of its binding sites in ESCs are not enriched for H3K9me3 (Fei et al., 2015). While SetDB1 was reported to interact with PRC2 to modulate H3K27me3 deposition, H3K27me3-modulation by SetDB1 appeared to be restricted to ∼5% of the target sites, as shown by the limited overlap between genomic loci bound by SetDB1, H3K27me3 and/or PRC2 (Fei et al., 2015). More recently, SetDB1 was shown to associate with Cohesin subunits SMC1A and SMC3 in ESCs, which are topological subunits of CTCF (Warrier et al., 2022). However, colocalization of SetDB1 and Cohesin was also observed in only ∼14% of the SetDB1 binding sites.

The protein domain composition of SetDB1 suggests the possibility of additional primary regulatory mechanisms beyond H3K9me3-, H3K27me3- and CTCF-mediated regulation. In addition to the SET domain, SetDB1 incorporates a methyl-CpG-binding domain (MBD). The MBD of SetDB1 has two arginine residues that enable DNA binding (Markouli et al., 2021). However, the MBD primarily appears to promote protein-protein interactions, including with DNA methyltransferases (DNMTs) that catalyze DNA methylation (DNAme) (Li et al., 2006; Markouli et al., 2021), another major heterochromatin mark. Li et al. (2006) demonstrated that SetDB1 directly interacts with DNMT3a and −3b, and showed the functional importance of this interaction through *in vitro* and *in vivo* studies. Regardless, the impact of loss of SetDB1 on DNAme through ESC differentiation has not been systematically explored.

While several individual regulatory mechanisms of SetDB1 have been characterized in ESCs, as outlined above, a comprehensive integration of these mechanisms and their mutual interplay is currently lacking. Given the fact that regions showing SetDB1-dependent H3K9me3 are frequently DNA methylated in serum ESCs, potential crosstalk with the DNAme machinery may be particularly important (Karimi et al., 2011; Leung et al., 2014; Matsui et al., 2010). In addition, H3K27me3 and DNAme are generally considered mutually exclusive (Brinkman et al., 2012; van Mierlo et al., 2019), and CTCF binding has been shown to be impaired by DNAme (Monteagudo-Sánchez et al., 2024). Given the central role of DNAme in the known regulatory modes of SetDB1, DNAme could be the missing link that integrates the other regulatory layers, potentially coordinating the complex regulatory network of SetDB1 during pluripotency.

Most studies on SetDB1 have been performed in serum-cultured mouse ESCs, which are hypermethylated. However, several recent studies have investigated the role of SetDB1 in naïve mouse ESCs (Wu et al., 2020; Fisher et al., 2017; Mochizuki et al., 2021; Deniz et al., 2018), which we previously have shown are hypomethylated (Habibi et al., 2013). While SetDB1 binding and H3K9me3 deposition appear similar in both ESC states, there are notable differences resulting from *Setdb1* knockout (KO). For example, in contrast to serum ESCs, SetDB1-deficient naïve ESCs undergo rapid cell death (Wu et al., 2020). Moreover, the activation of various SetDB1-targeted repeats following SetDB1 depletion is much more pronounced in naïve ESCs (Deniz et al., 2018). Taken together, these observations suggest that the underlying DNAme landscape may buffer against aberrant transcription of repeats that would normally be silenced at least in part by SetDB1 activity.

Therefore, we set out to comprehensively characterize the DNAme-mediated regulatory component of SetDB1 in pluripotent mouse ESCs through whole-genome bisulfite (WGBS)-based methylome time course analyses following *Setdb1* KO. Our analyses include both hypermethylated serum ESCs and hypomethylated naïve ESCs. By integrating the methylomes with DNA binding profiles of SetDB1, H3K9me3, H3K27me3, TET1, TET2, and CTCF, as well as RNA sequencing (RNA-seq)-based expression profiling of wild-type (WT) and *Setdb1* KO ESCs, we dissect the SetDB1-dependent DNAme mechanism and its integration with the other regulatory mechanisms across diverse genomic elements.

## Results

### Methylome profiling upon loss of SetDB1 reveals two distinct modes of action

To investigate the regulation of DNAme by SetDB1 during pluripotency, we made use of *Setdb1*^CKO/-^ conditional KO mouse ESCs, in which *Setdb1* KO can be induced upon administrating 4-hydroxytamoxifen (4-OHT) (Matsui et al., 2010) (Supplemental Fig. 1A). Induction with 4-OHT for 4 days facilitated the deletion of *Setdb1* floxed exons 15 and 16, which we validated by qPCR (Supplemental Fig. 1B), resulting in loss of SetDB1 as previously described (Matsui et al., 2010). To provide a global overview of DNAme levels upon SetDB1 depletion, we quantified DNAme through mass spectrometry (MS) at multiple time points following *Setdb1* KO in ESCs cultured in medium containing serum and LIF (abbreviated as ‘serum ESCs’) (Supplemental Fig. 1C). Similar to SetDB1-deficient primordial germ cells (PGCs) (Liu et al., 2014), we observed a substantial global increase of DNAme in serum ESCs upon induction of *Setdb1* KO. Global DNAme levels increased through 4 days of inducing *Setdb1* KO with 4-OHT, and peaked around day 10 post-KO induction. Extended culturing of SetDB1-deficient serum ESCs resulted in cell death around day 15 (Fig. 1A; Supplemental Fig. 1C).

**Figure 1:**
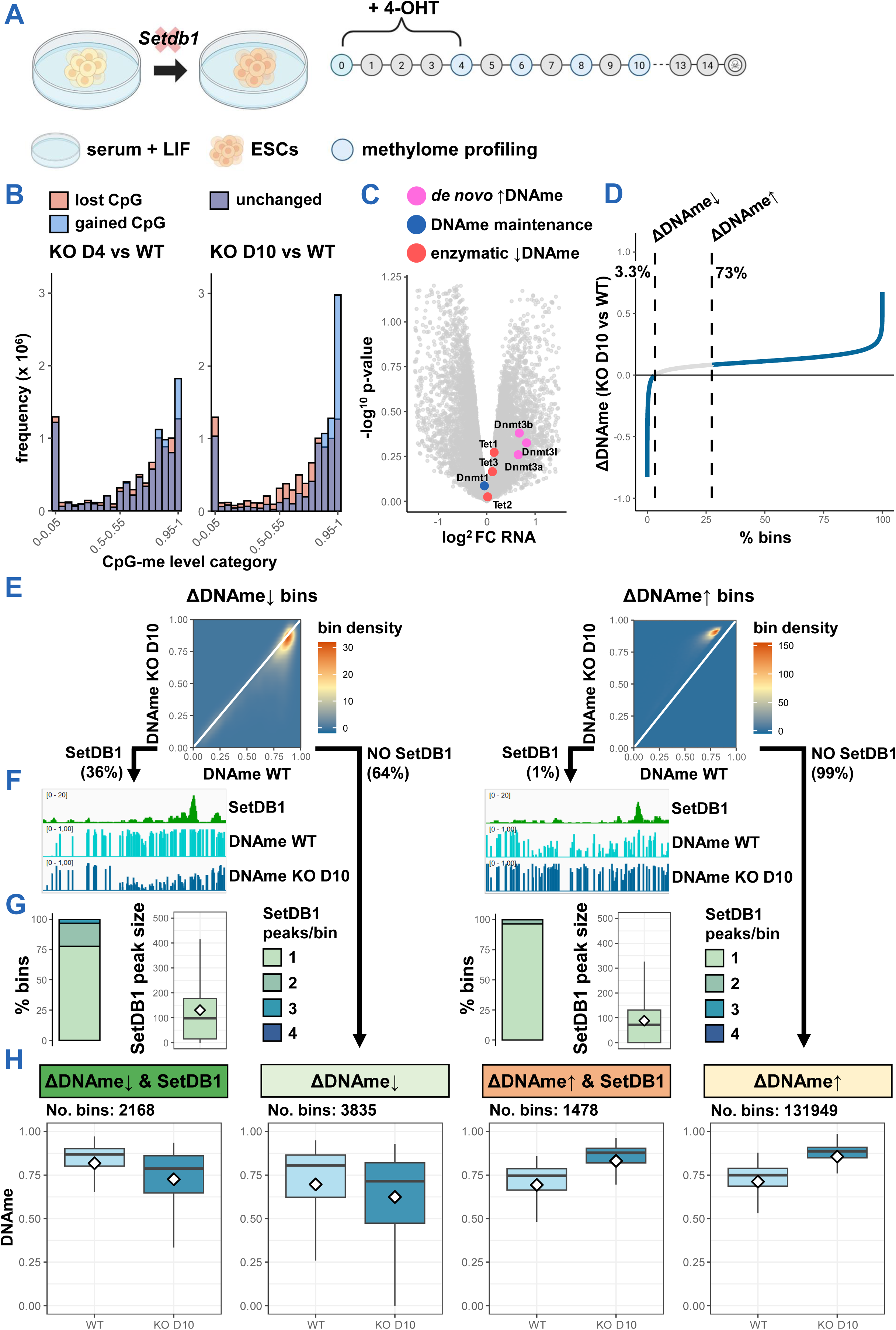
Methylome profiling in ESCs reveals that SetDB1 either directs DNAme or acts independently of it. **A**, Schematic of workflow for methylome profiling upon induction of *Setdb1* KO in serum ESCs. **B**, Bar graph of differential CpG methylation upon induction of *Setdb1* KO in serum ESCs. CpGs are categorized in aggregated methylation levels and sorted from lowest to highest on the horizontal axis. **C**, Volcano plot of RNA-seq-based differentially expressed genes in *Setdb1* KO serum ESCs compared to WT serum ESCs. Genes involved in the deposition and removal of DNAme are highlighted. **D**, Line graph illustrating methylome remodeling of 10kb genomic bins at 10 days post-*Setdb1*-KO induction in serum ESCs, with bins sorted by their average methylation change. Vertical dashed lines represent treshholds for ΔDNAme↓ and ΔDNAme↑ subsets. **E**, Heatmap showing methylation remodeling of bins previously categorized in D (6003 ΔDNAme↓ & 133,427 ΔDNAme↑ bins). Bin count density on color scale. Vertical white line represents slope = 1. **F**, IGV screenshots of SetDB1 ChIP-seq and WGBS data of exemplary ΔDNAme↓ and ΔDNAme↑ bins with SetDB1 binding. **G**, Integration of previously defined bins in D with nearby SetDB1 binding. SetDB1 peaks are allocated to ΔDNAme↓ bins and ΔDNAme↑ bins if within 1kb distance. Bar plot displays number of identified peaks near both bin categories. Boxplots represent a quantification of individual peak sizes (TMM-normalized values). White diamonds in boxplots depict mean TMM. **H**, Boxplots of DNAme in WT and *Setdb1* KO serum ESCs on day 10 at bins defined in D, further categorized by the presence or absence of SetDB1 peaks as defined in F. White diamonds represent mean DNAme.

Next, whole genome bisulfite sequencing (WGBS) was performed to examine the dynamic DNAme landscape at base pair resolution. In line with our MS data, the re-distribution of CpG methylation showed a global increase in DNAme after SetDB1 loss in serum ESCs (Fig. 1B; Supplemental Fig. 1D). A noticeable shift was detected in which the number of CpGs with low to moderate methylation levels (0-0.75) decreased, while those with high methylation levels (0.80-1.00) increased. Especially CpGs with moderate methylation levels progressively gained methylation to become hypermethylated by day 10 (Supplemental Fig. 2E). The global methylome expansion was evident across the entire genome, with CpGs on all chromosomes showing a gradual rise in methylation after *Setdb1* KO (Supplemental Fig. 1F). Potentially accounting for the genome-wide increase in DNAme, RNA-seq analyses of WT and *Setdb1* KO serum ESCs (Barral et al., 2022) revealed a >1.5-fold upregulation of *Dnmt3a* and *Dnmt3b de novo* DNMTs as well as their catalytically inactive stimulator *Dnmt3l* in *Setdb1* KO cells (Fig. 1C) (Suetake et al., 2004).

To further characterize differentially methylated regions in SetDB1-deficient serum ESCs and uncover the DNAme-mediated regulatory mechanisms of SetDB1, we divided the methylomes into 10kb bins. Ranking the bins by their average methylation change on day 10 revealed that 96.7% of bins showed an increase in methylation after *Setdb1* KO (Fig. 1D). To obtain a robust subset of bins with ΔDNAme↑ we omitted bins with minimal DNAme elevations relative to the global trend (the bottom 25th quantile; Fig. 1D). Opposing the global trend, a small group of demethylated bins (3.3%) was also observed, which we defined as ΔDNAme↓ regions. Notably, the demethylated ΔDNAme↓ bins were characterized by the highest endogenous DNAme levels in serum ESCs (Fig. 1E), suggesting a significant divergence from their default chromatin state after *Setdb1* KO.

To distinguish between direct and indirect effects of SetDB1 depletion on the methylome, we integrated our binned methylome data with SetDB1 chromatin immunoprecipitation sequencing (ChIP-seq) data (Warrier et al., 2022). SetDB1 peaks were detected within 1kb of both ΔDNAme↑ and ΔDNAme↓ bins, with comparable average peak sizes of SetDB1 (Fig. 1F-G). However, demethylated bins were targeted much more frequently by SetDB1 (36%) as compared to bins that gained methylation (1%). A similar trend was observed when examining individual CpGs, with their degree of demethylation correlating with increased levels of SetDB1 binding (Supplemental Fig. 1G). These results indicate that local demethylation was likely a direct result of proximal loss of SetDB1 binding. In contrast, SetDB1-targeted ΔDNAme↑ bins exhibited the lowest endogenous DNAme levels, and displayed a methylation increase after *Setdb1* KO that was similar to that of SetDB1-untargeted bins (Fig. 1H). Taken together, our findings suggest that SetDB1 operates through two distinct modes of action: (i) SetDB1 binds specific targets and guides the deposition of DNAme. Upon *Setdb1* KO, this DNAme enrichment is lost, possibly in part due to the absence of SetDB1-mediated guiding of the DNMT3 proteins (Li et al., 2006). (ii) At other regions, SetDB1 binds without influencing DNAme deposition. These target regions follow the global non-specific DNAme gain after *Setdb1* KO, which could be driven by increased and unguided activity of DNMT3A and/or DNMT3B, potentially independent of H3K9me3 deposition.

### Loss of SetDB1-mediated DNAme is tightly coupled to H3K9me3 erasure and pre-loaded TET2 near repeat elements

To determine whether targets with SetDB1-dependent DNAme coincide with H3K9me3 and H3K27me3 marks, we integrated the methylome binning with our previously generated H3K9me3 (Fisher et al., 2017) and H3K27me3 ChIP-seq data from Fei et al. (2015), generated from WT and *Setdb1* KO serum ESCs. Since these repressive histone marks have been demonstrated to cover large genomic regions (Chadwick & Willard, 2004; Pauler et al., 2009; Squazzo et al., 2006), we analyzed changes in H3K9me3 and H3K27me3 across the entirety of our previously defined methylome bins. This revealed that bins which gained DNAme post-KO were not enriched for H3K9me3, regardless of SetDB1 presence (Fig. 1H & 2A: orange categories of 1478 and 131,949 bins). Large H3K9me3 enrichments in WT serum ESCs were almost exclusively found on the ΔDNAme↓ SetDB1-targeted bins, where DNAme was also elevated (Fig. 1H & 2A: dark green category of 2168 bins). In addition to a reduction in DNAme after KO within these bins, H3K9me3 was erased to levels comparable to those of other bins. These results indicate that the regulatory modes of SetDB1 through DNAme and H3K9me3 are tightly coupled. Conversely, only minor changes in H3K27me3 deposition were observed after *Setdb1* KO among all bin categories (Fig. 2B).

**Figure 2:**
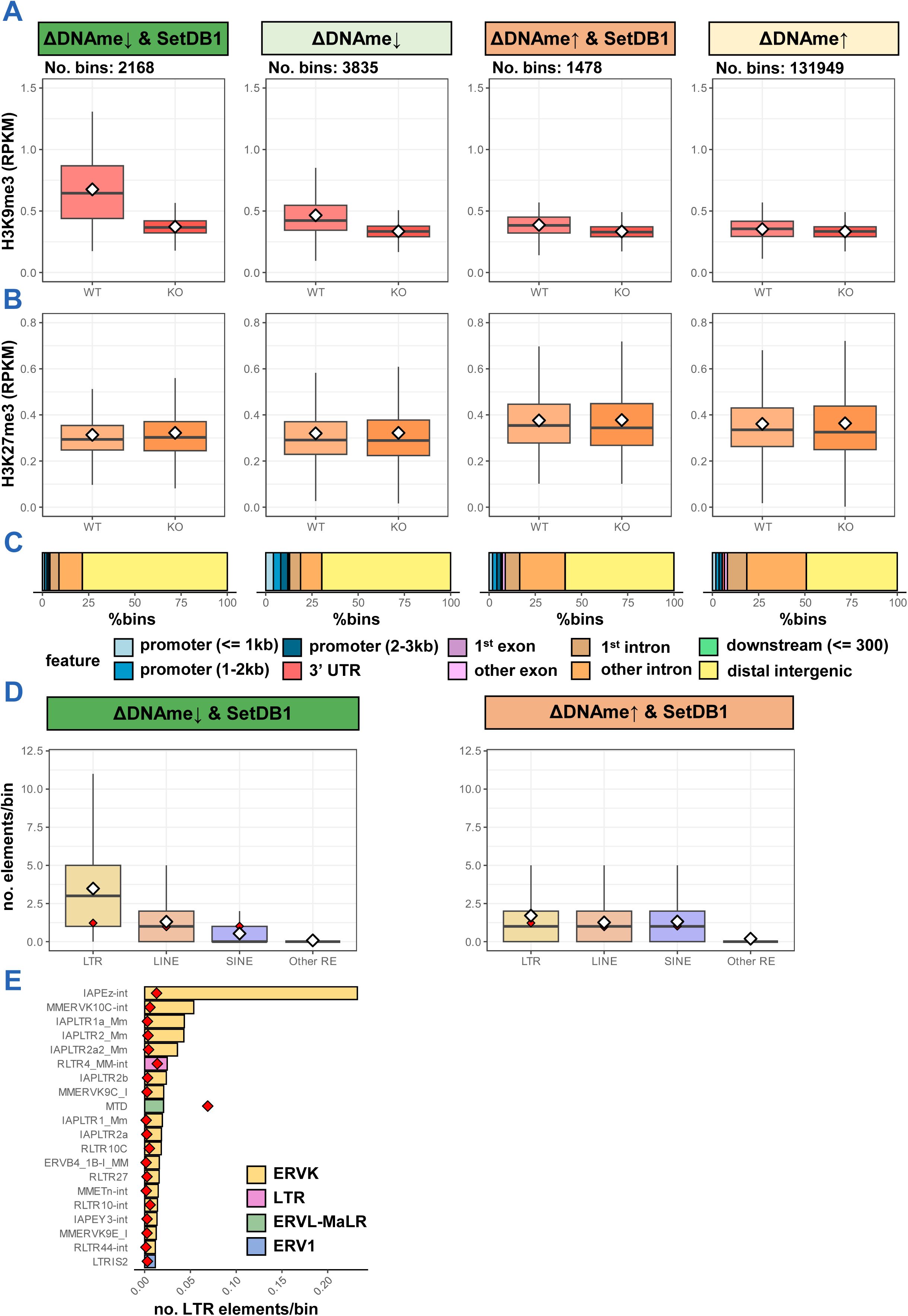
SetDB1-mediated DNAme and SetDB1-catalyzed H3K9me3 are tightly coupled. **A**, Boxplots of H3K9me3 levels over the entirety of methylome bins, as defined in 1H, in WT and *Setdb1* KO serum ESCs. White diamonds represent averages. **B**, Boxplots of H3K27me3 levels, as in A. **C**, Bar plot showing the overlapping genomic features at methylome bin centers, as grouped in A. Genomic annotations determined by ChIPseeker. **D**, Boxplots showing the number of overlapping repeat elements per methylome bin from the SetDB1-targeted ΔDNAme↓ and ΔDNAme↑ bin groups. The other RE group includes scRNAs, rRNAs, snRNAs, srpRNAs, DNA repeats, satellites, RCs, Retroposons and tRNAs. Expected averages in any 10kb bin plotted as red diamonds. **E**, Bar graph of observed and expected frequency (red diamonds) of the 20-most-occurring LTRs in SetDB1-targeted ΔDNAme↓ bins.

Next, we set out to understand what genomic features are associated with SetDB1 targets that lose both H3K9me3 and DNAme after *Setdb1* KO. SetDB1-targeted ΔDNAme↓ bins were predominantly located in distal intergenic regions, with only few associations to genes (Fig. 2C). Comparatively, SetDB1-associated ΔDNAme↑ bins were enriched with genic features, especially intronic elements. This suggests DNAme-H3K9me3-mediated regulation by SetDB1 may be associated with non-coding and repetitive elements, consistent with the proposed primary role of SetDB1 in mouse ESCs (Karimi et al., 2011; Matsui et al., 2010). To further refine our analysis, we determined the numbers of overlapping retrotransposons from the LTR, LINE, SINE, and other repeat families (Other RE) per bin, and compared that with expected counts based on random distribution (red diamonds) (Fig. 2D). The SetDB1-targeted ΔDNAme↓ bins are highly enriched for LTR retrotransposons, in particular from the ERVK family, including IAPEz elements (17.9x enrichment) (Fig. 2E). This aligns with previous studies showing that regulation members of the IAPEz subfamily involves SetDB1-dependent H3K9me3 and DNAme (Karimi et al., 2011; Leung et al., 2014). Other enriched ERVKs in SetDB1-targeted ΔDNAme↓ bins include other IAP (9.6x enrichment), MMERVK (7.4x enrichment) and RLTR (4.5x enrichment) subfamilies (Fig. 2E). In contrast, SetDB1-targeted ΔDNAme↑ bins are not enriched for LTRs, nor any other repeat element (Fig. 2D).

Leung et al., (2014) proposed that the SetDB1-H3K9me3 axis protects against demethylation, aligning with our observation that bins marked with SetDB1-dependent H3K9me3 show reduced DNAme after *Setdb1* KO. However, we noticed that changes in H3K9me3 and DNAme levels did not occur simultaneously on a global level: While SetDB1-dependent H3K9me3 is effectively lost within 4 days of inducing *Setdb1* KO (Fisher et al., 2017; Matsui et al., 2010), changes in DNAme peak 10 days post-KO induction (Fig. 1B). To investigate the functional relationship between DNAme and H3K9me3 at SetDB1 targets over time, we made use of our WGBS time course to assess their dynamics following *Setdb1* KO. As such, we defined fast and slow methylome bins based on whether the greatest absolute DNAme change was observed early (day 4 or 6) or late (day 8 or 10) (Supplemental Fig. 2A). Demethylated SetDB1-target bins were clearly distinguished by either an immediate or a delayed removal of DNAme (Supplemental Fig. 2B). The slowly demethylated target bins further differed by modestly higher levels of DNAme, both at the start and at day 10 post-KO induction. The demethylation dynamics of these slow bins seem to match the dynamics of passive dilution of DNAme, likely due to failure of DNMT3 recruitment and/or failure of DNMT1 recruitment to H3K9me3-marked regions (Fang et al., 2016). Of note, in contrast to fast and slow demethylation, fast and slow DNAme gains were not clearly separated (Supplemental Fig. 2B).

To assess whether the demethylation rate after SetDB1 loss is dependent on H3K9me3 dynamics, we assayed H3K9me3 patterns among fast and slowly demethylated SetBD1 target bins. Interestingly, we found no apparent difference in endogenous H3K9me3 levels between fast and slowly demethylated target bins (Supplemental Fig. 2C-D). Also, both bins show complete erasure of H3K9me3 after 6 days of SetDB1 loss (Supplemental Fig. 2C-D) (Fisher et al., 2017). Furthermore, H3K27me3 levels are low/absent in the SetDB1-targeted ΔDNAme↓ bins (Fig. 2B) and we did not observe global differences in H3K27me3 in WT cells nor in its remodeling after *Setdb1* KO when comparing fast and slowly demethylated target bins (Supplemental Fig. 2C-D).

Notably, a range MMERVKs, RLTRs and (MM-)ETn copies were primarily found in rapidly demethylated bins, whereas IAPs, including IAPEz elements, were mostly located in slowly demethylated bins (Supplemental Fig. 2E). In recent years, the family of TET proteins has been identified as potent DNA demethylation enzymes (e.g. Deniz et al., 2018; Lio et al., 2020; de la Rica et al., 2016). As such, we hypothesized that the fast demethylated bins lose DNAme through passive dilution and additional rapid removal of DNAme by pre-loaded TETs (Leung et al., 2014). To test this hypothesis, we utilized previously generated TET1 and TET2 ChIP-seq profiles from serum ESCs (Xiong et al., 2016; Williams et al., 2011), and plotted these over the various ERVKs that are enriched in the fast and slowly demethylated SetDB1 target bins (Supplemental Fig. 2F). TET1 targeting was uniform and did not exhibit a clear preference for either fast or slowly demethylated ERVKs. On the other hand, fast-demethylated ERVKs were distinguished by high TET2 pre-loading in serum ESCs, while slowly demethylated ERVKs were not endogenously targeted by TET2. To evaluate the enzymatic activity of pre-loaded TET enzymes, we mined profiles of TET-mediated oxidation products (5hmC, 5fC, and 5caC) upon *Tdg* knockdown in serum ESCs (Shen et al., 2013). Depletion of TDG leads to the accumulation of TET-mediated oxidation products by blocking the final step of active demethylation. Following *Tdg* knockdown in serum ESCs, many ERVK subfamilies from the fast ΔDNAme↓ category exhibit elevated levels of 5caC, the terminal TET-mediated oxidation product, indicating active DNAme turnover (Supplemental Fig. 2G). In contrast, ERVK subfamilies from the slow ΔDNAme↓ category are not enriched with 5cAC after *Tdg* knockdown, but accumulate 5fC instead. This suggests a lack of complete, active DNAme turnover at the slowly demethylated ERVK subfamilies. Together, our results show that specific ERVK subfamilies, such as MMERVK9s, are pre-loaded with TET2 and display active DNAme turnover, and rapid demethylation following SetDB1 loss. Other elements, particularly IAPs, might be shielded from active demethylation under WT conditions by protection against TET2 pre-loading. Protection against rapid demethylation of IAP elements may be particularly important, as IAPs are still actively retrotransposing in the rodent lineage and are in turn highly mutagenic in both germline and somatic lineages (Maksakova et al., 2006).

### SetDB1 loss triggers DNAme-H3K9me3 erasure and reactivation of targeted ERVKs and nearby genes, and ICRs

Our binning approach does not easily allow the investigation of transcriptional responses from individual genomic elements. To facilitate such analysis, we next projected the data on the genomic loci directly. Consistent with earlier results (Fig. 1 & 2), various ERVK subfamilies are targeted by SetDB1 and enriched with DNAme and H3K9me3 (Supplemental Fig. 3A). Among these, IAP elements show the greatest SetDB1 targeting and enrichment of DNAme and H3K9me3. A minority of LTR regions of IAPs (generally referred to as IAP-LTR) and other ERVKs are also marked with H3K27me3. A similar co-occurrence of H3K9me3 and H3K27me3 was also observed in PGCs, including at hypermethylated IAPs (Liu et al., 2014). However, H3K27me3 coverage is generally greatest on ERVKs and other LTRs that lack H3K9me3 (and DNAme) enrichment in serum ESCs (Supplemental Fig. 3B). This aligns with previous findings that H3K9me3 and H3K27me3 mostly mark distinct genomic regions in ESCs (Fei et al., 2015), and supports the hypothesis that H3K27me3 may repress ERVs that are not marked by SetDB1-dependent H3K9me3 and DNAme (Prokopuk et al., 2017). Notably, H3K27me3-enriched LTRs are rarely bound by SetDB1 compared to DNAme-H3K9me3-enriched LTRs (Supplemental Fig. 3A), suggesting that SetDB1 preferentially regulates its retrotransposon targets via H3K9me3 in combination with DNAme. Furthermore, consistent with Figures 1 and 2, strongly targeted ERVKs were demethylated and completely lost their H3K9me3 enrichment after *Setdb1* KO (Fig. 3A; Supplemental Fig. 3C-D). Following the loss of DNAme and H3K9me3 on ERVKs, we noticed a minor increase in H3K27me3 (Fig. 3A), which is in line with our previous observation that loss of DNAme allows PRC2 to deposit diffuse H3K27me3 (Van Mierlo et al., 2019; Walter et al., 2016).

**Figure 3:**
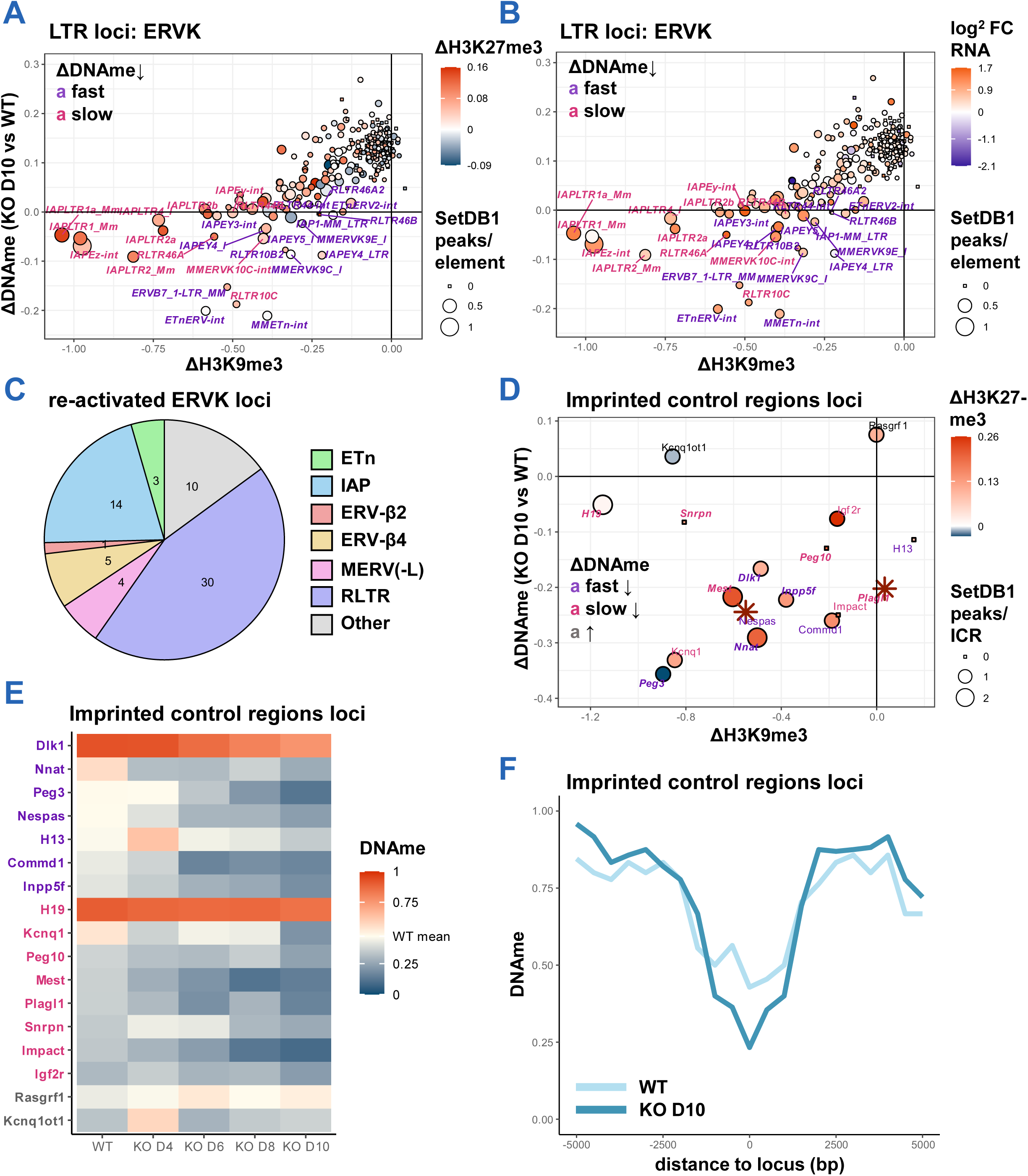
ERVKs and ICRs with SetDB1-dependent DNAme-H3K9me3 are re-activated in serum ESCs following SetDB1 loss. **A**, Scatter plot of LTR-ERVKs showing ΔH3K9me3 (ΔRPKM), ΔDNAme (KO day 10 vs WT), and ΔH3K27me3 (ΔRPKM) of ERVKs in serum ESCs upon *Setdb1* KO. Number of SetDB1 ChIP-seq peaks per of ERVK copy (1kb proximity) displayed as point size. **B**, Scatter plot similar to A, but color scale now represents difference of expression after *Setdb1* KO, as log^2^ fold change. **C**, Pie chart illustrating the number of increasingly expressed ERVKs upon *Setdb1* KO in serum ESCs from different ERVK-groups (fold-change > 1.5). **D**, Scatter plot of ICRs, similar to A. Names in italic font if imprinted gene was transcriptionally upregulated after *Setdb1* KO. **E**, Heatmap of DNAme levels at ICRs in WT and *Setdb1* KO serum ESCs. ICRs are organized by label colors assigned in D. **F**, Line graph illustrating median methylation of ICRs (zero distance) and their immediate surrounding genomic.

To assay whether the chromatin remodeling at ERVKs following *Setdb1* KO results in their reactivation, we re-analyzed RNA-seq data generated from serum ESCs after *Setdb1* KO (Barral et al., 2022). Over 80 percent of ERVK subfamilies show an increase in transcription after *Setdb1* KO (Fig. 3B). SetDB1-targeted ERVKs that lost DNAme and H3K9me3 post-KO were upregulated by up to 3.3-fold. This indicates that the minor increase of H3K27me3 after SetDB1 depletion (Fig. 3A) is insufficient to keep these retrotransposons repressed. Notably, upregulated ERVKs (fold change > 1.5) include a large range of IAP, ETn, ERV-β2/4 and MERV elements (Fig. 3C), all of which are capable of retrotransposition and pose a threat to genome integrity through insertional mutagenesis (Gagnier et al., 2019; Maksakova et al., 2006). These findings align with the previously proposed model that SetDB1 acts as a guardian of the genome by protecting against activation of parasitic elements during embryogenesis (Matsui et al., 2010; Leung & Lorincz, 2012).

The presence of both DNAme and H3K9me3 is not exclusive to LTR-ERVKs, as this combination has also been observed at imprinted control regions (ICRs) (Fisher et al., 2017; Habibi et al., 2013; Leung et al., 2014), suggesting their potential dependency on SetDB1-mediated regulation. Indeed, 13 out of 17 (76%) ICRs described in the literature (Betto et al., 2021; Santini et al., 2021) are targeted by SetDB1, as indicated by 1-2 SetDB1 peaks within 1kb distance (Fig. 3D). The DNAme level across the ICR averaged 47% and many ICRs were densely marked with H3K9me3 (Supplemental Fig. 3E). After *Setdb1* KO in serum ESCs, most ICRs lost H3K9me3 as well as proximal DNAme, the epigenetic mark essential for classical imprinting (Fig. 3D-F; Supplemental Fig. 3F-G). Some ICRs were more susceptible to demethylation than others (Supplemental Fig. 3F-G), akin to specific ERVKs. Deposition of H3K27me3 slightly increased following *Setdb1* KO (except on the *Kcnq1ot1* ICR), but this was insufficient to prevent the de-repression of several imprinted genes (Fig. 3D; italic names). Overall, our findings highlight a strong resemblance between SetDB1-mediated regulation of ICRs and ERVKs through a repressive DNAme-H3K9me3 axis in serum ESCs.

Our binning approach revealed that the SetDB1-targeted ΔDNAme↓ regions also include a small number of promoters and other genic elements (Fig. 2C). To examine which genes are regulated by the DNAme-mediated component of SetDB1, we investigated SetDB1-targeted genic loci. We observed only 37 demethylated target genes (ΔDNAme < −0.05) within 1kb proximity of at least one SetDB1 ChIP-seq peak. Similar to ERVKs and ICRs, demethylated target genes also lost their endogenous H3K9me3 enrichment after *Setdb1* KO (Fig. 4A). Following *Setdb1* KO, 78 percent of demethylated target genes were increasingly expressed, up to 2.6-fold (i.e. *Sult2a3*) (Fig. 4B). GO-enrichment analyses revealed enriched terms related to meiosis (Fig. 4C), consistent with previously reported functions of DNAme-H3K9me3-regulated target genes of SetDB1 (Karimi et al., 2011; Leung et al., 2014). To find out whether nearby ERVKs are involded in the regulation of these genes, we probed the number of demethylated ERVKs within 5kb proximity of demethylated target genes (Fig. 4D). An enrichment for demethylated ERVKs (expected: 0.08 demethylated ERVKs per gene, red diamond) was found near demethylated target genes, showing more than two nearby demethylated ERVKs on average. Several of these ERVKs are embedded within gene regulatory elements, such as promoters, though the majority is found in intronic regions (Fig. 4E). For example, the derepressed *Zfp951* gene, that lost DNAme and H3K9me3 after *Setdb1* KO, embeddes six demethylated ERVKs near its locus (Fig. 4F). Hence, the DNAme-H3K9me3-mediated repression of a majority of these genes may have originated from the targeting and regulation of flanking ERVKs by SetDB1. Notably, Karimi et al. (2011) reported that aberrant re-activation of ERVKs upon *Setdb1* KO is associated with disturbed expression of neighboring genes. Taken together, these observations suggest that ERVKs exert a DNAme- and H3K9me3-dependent repressive influence on nearby target genes, a mechanism that is disrupted upon SetDB1 depletion.

**Figure 4:**
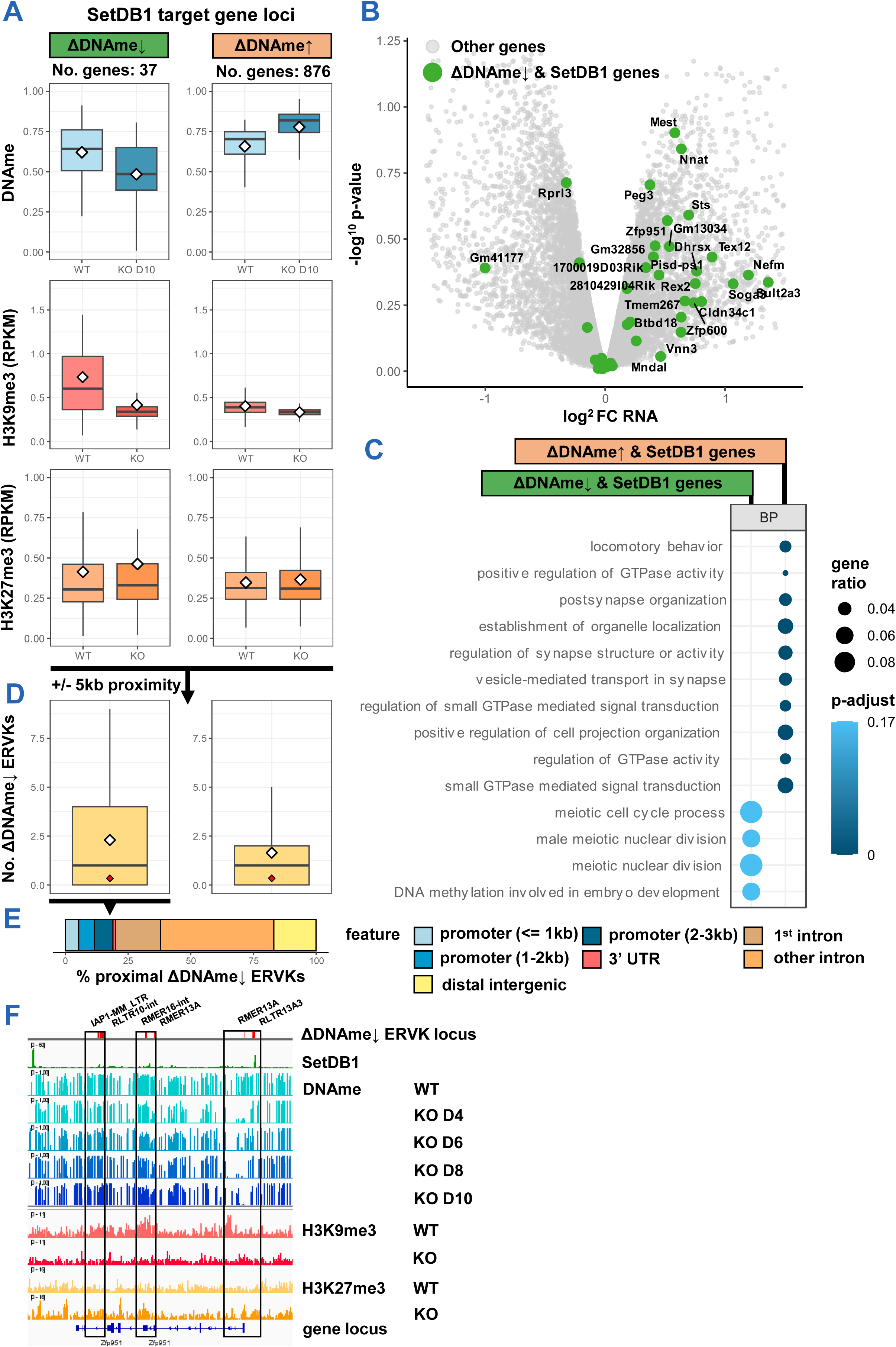
Genes with SetDB1-dependent DNAme-H3K9me3 are de-repressed in serum ESCs following SetDB1 loss. **A**, Boxplots of DNAme, H3K9me3, and H3K27me3 levels of SetDB1-targeted genes in WT and *Setdb1* KO serum ESCs, grouped by methylation change at day 10 of *Setdb1* KO. **B**, Volcano plot of differentially expressed genes in *Setdb1* KO serum ESCs compared to WT serum ESCs, as observed in RNA-seq data. Demethylated SetDB1-target genes are highlighted in green. **C**, Dot plot showing biological processes enriched among SetDB1-targeted genes (demethylated or increasingly methylated by day 10 of *Setdb1* KO), as determined by GO-analysis. **D**, Boxplots showing number of demethylated ERVK copies within 5kb distance of SetDB1-target genes, as categorized in G. White diamonds represent averages. Red diamonds represent expected number of demethylated ERVKs near any gene. **E**, Barplot showing genomic positions of demethylated ERVK elements near demethylated SetDB1 target genes. Genomic annotations determined by ChIPseeker. **F**, IGV screenshots of the *Zfp951* locus. Demethylated ERVK (by day 10 of *Setdb1* KO) loci indicated in red above IGV tracks.

### H3K27me3- and DNAme-H3K9me3-mediated regulatory mechanisms of SetDB1 are uncoupled

A substantial fraction of SetDB1-targeted bins does not contain SetDB1-dependent DNAme and H3K9me3, and displays DNAme increases following *Setdb1* KO that mirror the global, non-specific trend (Fig. 1 & 2: ΔDNAme↑ & SetDB1 bin category). Here, we examine what regulatory role SetDB1 has at these sites. Fei et al. (2015) demonstrated that a subset of SetDB1 target regions does not contain H3K9me3, but overlap with repressive H3K27me3 marks instead, regulating nearby neural development genes. Therefore, we asked whether the SetDB1-targeted bins without SetDB1-dependent DNAme represent these H3K27me3-marked loci. To enable direct comparison with our data, we conducted a SetDB1 peak-centered analysis. To define SetDB1 target regions with and without SetDB1-dependent DNAme in this analysis, we ranked all 10kb regions surrounding a SetDB1 peak summit based on their DNAme changes after *Setdb1* KO. We selected fractions above the 25th percentile of both demethylated and increasingly methylated groups to obtain subsets with and without SetDB1-dependent DNAme, respectively (Fig. 5A: ΔDNAme↓, 21% and ΔDNAme↑, 29%). Of note, this peak centered analysis confirmed our previous observations. Reassessment of DNAme and H3K9me3 levels across these peak regions confirmed the tight coupling between DNAme and H3K9me3 enrichments under WT conditions, and their erasure after *Setdb1* KO (Fig. 5A; Supplemental Fig. 4A-B).

**Figure 5:**
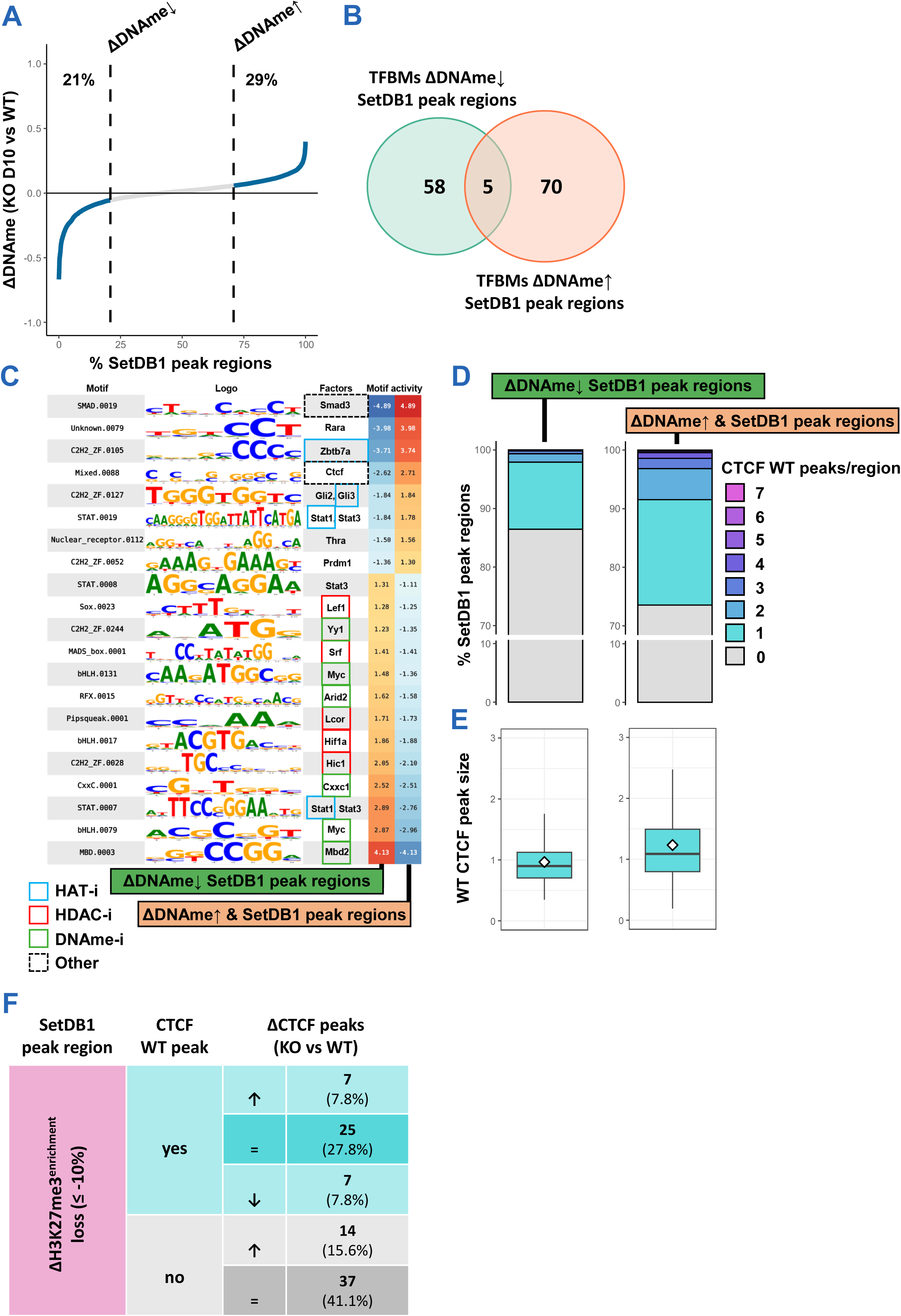
Targets without SetDB1-mediated repression are enriched with binding motifs of CTCF, SMAD3 and other TFs that allow regulation beyond heterochromatin formation. **A**, Line graph illustrating methylome remodeling of SetDB1 peak regions (10kb centered on ChIP-seq peak summits) at 10 days post-*Setdb1* KO induction in serum ESCs. Vertical dashed lines represent 25^th^ quantiles of demethylated and increasingly methylated peak regions. **B**, Venn diagram showing overlap between TFBMs that are enriched (according to Gimme Maelstrom) in SetDB1 binding sites (200bp centered on ChIP-seq peak summit) within categories described in A. **C**, Schematic representation that highlights selection of enriched TFBMs in SetDB1 peak region categories as defined in A. **D**, Bar graph showing number of CTCF ChIP-seq peaks from WT serum ESCs near SetDB1 peak regions (within 1kb distance), as categorized in A. **E**, Boxplots showing sizes of individual CTCF peaks in WT serum ESCs, as read pileups normalized to 10bp, calculated by Macs2, near SetDB1 peak region categories from A. **F**, Table summarizing CTCF peak remodeling at H3K27me3-mediated SetDB1 peak regions. Numbers reflect number and percentage of H3K27me3-mediated SetDB1 peak regions within categories.

Plotting the H3K27me3 genomic profiles (Fei et al., 2015) over the SetDB1 peak regions confirmed that approximately 5% of SetDB1 peak regions exhibited endogenous H3K27me3 enrichment (Supplemental Fig. 4C). Remapping the H3K27me3 enrichments to the ΔDNAme-ranking of SetDB1 peak regions (Supplemental Fig. 4D-E) revealed that most H3K27me3 enrichments were located on SetDB1 peak regions that gained DNAme post-*Setdb1* KO (Supplemental Fig. 4F: ΔDNAme↑ category), which endogenously contained low levels of DNAme (Supplemental Fig. 4A) and H3K9me3 (Supplemental Fig. 4B). As such, almost all of these H3K27me3-decorated regions do not overlap with SetDB1-targeted ΔDNAme↓ bins (Fig. 1-2). Notably, only 22.8% of H3K27me3-enriched SetDB1 peak regions (equivalent to 1.14% of total SetDB1 peak regions) experienced a loss of 10% or more of their H3K27me3 coverage following *Setdb1* KO (Supplemental Fig. 4C-F: dark purple category), representing targets regulated through SetDB1-mediated H3K27me3. These SetDB1-mediated H3K27me3 targets are almost entirely confined to ΔDNAme↑ SetDB1 peak regions (Supplemental Fig. 4D & 4F), and show relatively strong localization towards promoter and exonic regions of genes (Supplemental Fig. 4G). Together, these findings indicate that the H3K27me3- and DNAme-H3K9me3-mediated regulatory mechanisms of SetDB1 are uncoupled.

### ΔDNAme↑ SetDB1 binding sites are enriched with TF motifs that allow regulation beyond heterochromatin formation

Considering the limited extent of SetDB1-mediated H3K27me3 regulation among target regions without SetDB1-mediated DNAme, we explored what other SetDB1 regulatory mechanisms might be operating at these sites. Thereto, we analyzed and compared transcription factor binding motifs (TFBMs) among the binding sites of SetDB1. We extracted the nucleotide sequences of SetDB1 binding sites (i.e. 200bp surrounding a peak summit) from peak regions that showed ΔDNAme↑ or ΔDNAme↓ post-KO (Fig. 5A), and identified enriched TFBMs using Gimme Maelstrom (Bruse & Heeringen, 2018). We identified 75 enriched TFBMs within ΔDNAme↑ SetDB1 binding sites, 70 of which were not enriched among SetDB1 ΔDNAme↓ binding sites (Fig. 5B).

TFBMs unique to ΔDNAme↑ SetDB1 binding sites were linked to transcription factors associated with histone acetyltransferase binding and transcriptional activation (Fig. 5C; Supplemental Fig. H), indicative of SetDB1 recruitment to non-heterochromatic regions. We also observe a strong enrichment of SMAD3 and CTCF motifs here (Fig. 5C). SMAD3 is the primary mediator of TGF-β signaling, and the SMAD3-SetDB1 axis has previously been implicated in epithelial-mesenchymal transition (EMT) in disease states (Du et al., 2018; Liu et al., 2022). In a similar manner, the SMAD3-SetDB1 identified here could influence the EMT during gastrulation. The co-occurrence of SetDB1 binding and CTCF motifs suggests that SetDB1 may contribute to large-scale 3D chromatin organization (Tam et al., 2024). Interestingly, CTCF binding is known to be highly sensitive to DNAme levels at its binding sites and surrounding regions, with DNAme generally impairing CTCF binding and chromatin looping (Monteagudo-Sánchez et al., 2024). This aligns with our observation that CTCF motifs are particularly enriched among SetDB1 binding sites in peak regions that contain relatively low DNAme under WT conditions (Supplemental Fig. 4A: ΔDNAme↑ category; cf. ΔDNAme↓ category). This observation parallels the results of Tam et al. (2024), who reported that SetDB1-dependent H3K9me3 – tightly coupled to DNAme in our data – and CTCF are mutually exclusive.

To further substantiate the enrichment of CTCF motifs among the ΔDNAme↑ SetDB1-target regions, we overlayed the SetDB1 peak regions with CTCF ChIP-seq profiles of WT and *Setdb1* KO serum ESCs (Tam et al., 2024). This revealed that 2.7 times more ΔDNAme↑ SetDB1 peak regions overlapped with CTCF than ΔDNAme↓ SetDB1 peak regions (Fig. 5D; 619/2340 = 26% compared to 227/1676 = 13%). Endogenous CTCF peaks among ΔDNAme↑ SetDB1 peak regions were also slightly larger on average, and contained more pronounced outliers (Fig. 5E), consistent with the previously reported inhibiting effect of DNAme on CTCF binding (Monteagudo-Sánchez et al., 2024). Following SetDB1 depletion, both ΔDNAme↓ and ΔDNAme↑ SetDB1 peak regions gained CTCF binding (Supplemental Fig. 4I). However, CTCF peak sizes especially increased in ΔDNAme↓ SetDB1 peak regions after *Setdb1* KO, potentially due to the removal of DNAme that previously obstructed endogenous CTCF binding at these sites. Likewise, SetDB1 peak regions that lacked endogenous CTCF and underwent demethylation after *Setdb1* KO, gained the most CTCF peaks among all SetDB1 target sites post-KO (Supplemental Fig. 4J).

To assess whether CTCF modulation by SetDB1 operates independently of its H3K27me3 regulatory mechanism, we assayed the number of CTCF peaks before and after *Setdb1* KO at H3K27me3-mediated targets. Most of these targets were not endogenously bound by CTCF (Fig. 5F), and did not acquire CTCF binding following *Setdb1* KO. Although more than a quarter of H3K27me3-mediated SetDB1 targets was natively associated with CTCF, CTCF peak numbers remained unchanged here after SetDB1 loss, suggesting that CTCF regulation occurred independently of SetDB1. Collectively, our analyses revealed two uncoupled repressive mechanisms of SetDB1. Most repressed targets are regulated through a DNAme-H3K9me3 dual silencing mechanism, whereas a minor subset is regulated through H3K27me3. In addition, SetDB1 modulates CTCF binding at a subset of target sites independently of its repressive functions. Notably, non-repressed SetDB1 targets are also highly enriched for SMAD3 and other TF motifs, highlighting a broader role of SetDB1 in genome regulation beyond heterochromatin formation.

### Loss of SetDB1-dependent DNAme reveals repressive buffering deficit in ground-state ESCs

The co-occupation of SetDB1-mediated H3K9me3 and DNAme raises the question what regulatory role DNAme fulfills at repressed targets - specifically, why DNAme is always coupled to the catalytic deposition of H3K9me3 by SetDB1 in serum ESCs. While serum ESCs exhibit a hypermethylated metastable state, the hypomethylated ground-state of pluripotency can be modeled *in vitro* by ESCs cultured in the presence of two inhibitors for MEK and GSK-3, along with LIF (referred to as ‘2i ESCs’). These 2i ESCs provide a powerful platform for investigating the regulatory mechanisms of SetDB1 within a chromatin state that is largely devoid of DNAme. Therefore, we assessed the functional importance of DNAme by extending our integrative methylome analyses to include both WT and *Setdb1* KO 2i ESCs.

In line with previous observations (Wu et al., 2020), SetDB1 depletion upon addition of 4-OHT to *Setdb1*^CKO/-^ 2i ESCs (Supplemental Fig. 1B) resulted in reduced cell growth, followed by massive cell death at day 4-5 (Fig. 6A; Supplemental Fig. 1C). As expected, the chromatin landscape of 2i ESCs is largely demethylated, which persisted upon SetDB1 depletion (Fig. 6B; Supplemental Fig. 5A-C). We observed a slight, global increase in DNAme according to MS (Supplemental Fig. 1C) and WGBS (Fig. 6B; Supplemental Fig. 5A-C), but this was minor compared to the increase observed in SetDB1-depleted serum ESCs (Fig. 1B; Supplemental Fig. 1D-F). The ΔDNAme↓ SetDB1 targets as defined in serum ESCs showed slightly elevated DNAme levels in 2i ESCs, but at much lower levels compared to serum ESCs (Fig. 6C-D). Notably, we did not detect any substantial DNAme remodeling at these target regions in 2i ESCs following *Setdb1* KO, not even in areas that underwent the most extensive remodeling in serum ESCs (Fig. 6C). This suggests that the small DNAme deposits in 2i ESCs are maintained independently of SetDB1, at least within the 4-day time frame of *Setdb1* KO. Overall, DNAme-mediated regulation at SetDB1 target sites was largely absent in 2i ESCs compared to serum ESCs.

**Figure 6:**
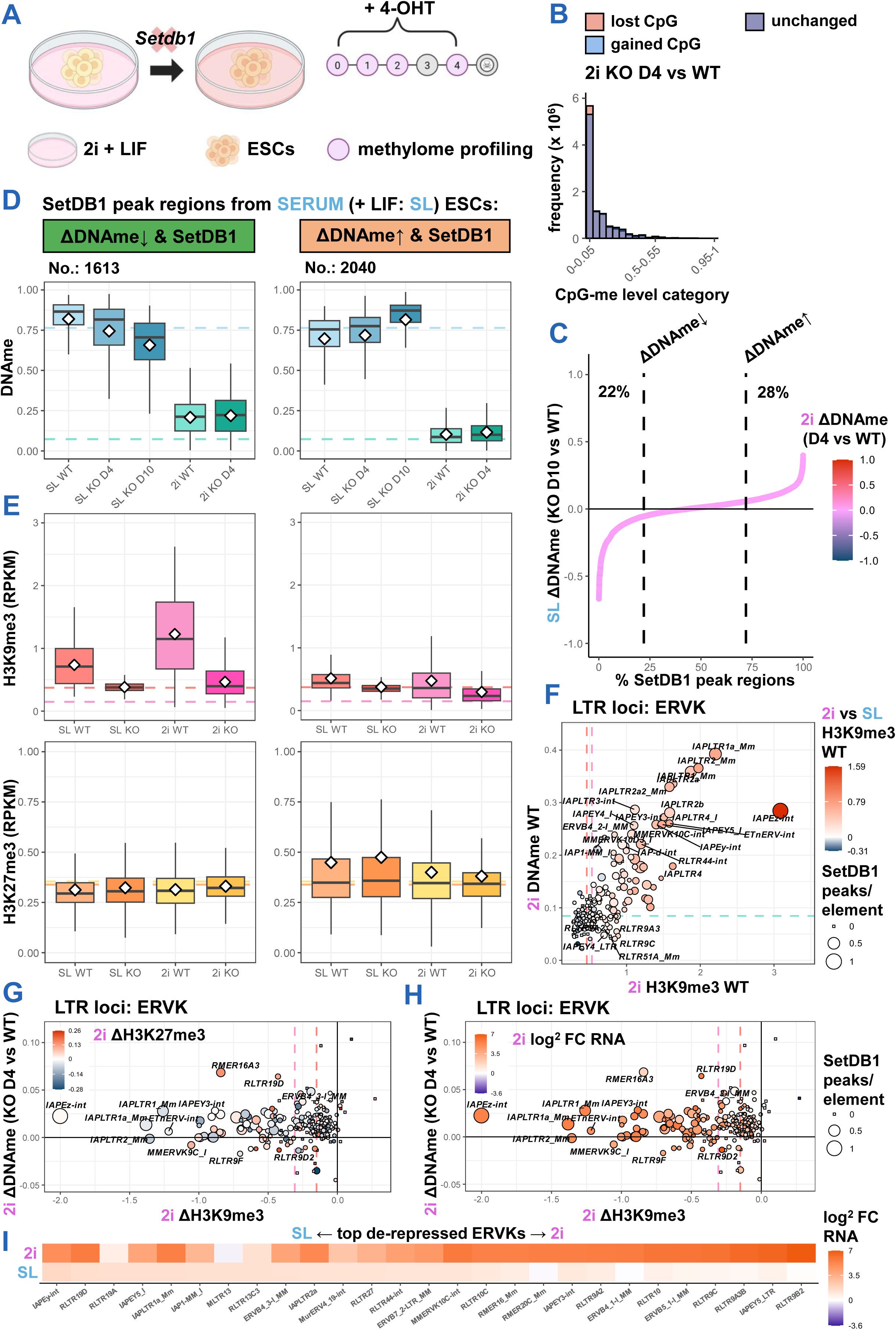
Hypomethylated naive ESCs strongly depend on SetDB1-mediated H3K9me3 to repress ERVKs, leaving these repeats prone to a high level of re-activation upon *Setdb1* KO. A, Schematic of workflow for methylome profiling upon induction of *Setdb1* KO in 2i ESCs. **B**, Bar graph of differential CpG methylation upon *Setdb1* KO in 2i ESCs, similar to Fig. 1B. **C**, Line graph, similar to Fig. 5A, but dot colors reflect DNAme change observed upon induction of *Setdb1* KO in 2i ESCs. **D**, Boxplots of DNAme in WT and *Setdb1* KO serum and 2i ESCs at target region categories defined in Fig. 5A. Dotted lines reflect median WT DNAme levels in serum ESCs (light blue) and 2i ESCs (turquois). **E**, Boxplots of H3K9me3 and H3K27me3 in WT and KO serum and 2i ESCs at target region categories defined in D. Dotted lines reflect median WT levels of H3K9me3 (red for serum ESCs, pink for 2i ESCs) and H3K27me3 (orange for serum ESCs, yellow for 2i ESCs). **F**, Scatter plot of ERVKs in 2i ESCs showing WT DNAme and WT H3K9me3 levels, and the difference in WT H3K9me3 between 2i and serum ESCs (colour scale). The number of SetDB1 ChIP-seq peaks per of ERVK copy in serum ESCs (within 1kb proximity) are displayed as point size. Labels are carried over from Fig. 3A. Dotted lines represent averages of corresponding axis parameters (WT H3K9me3: red for serum ESCs, pink for 2i ESCs and WT DNAme: turquois for 2i ESCs). **G**, Scatter plot of ERVKs in 2i ESCs upon induction of *Setdb1* KO, similar to Fig. 3A. Labels are carried over from Fig. 3A. Dotted lines represent averages of corresponding axis parameters (red for serum ESCs, pink for 2i ESCs). **H**, Scatter plot similar to G, but color scale now represents the difference in expression upon induction of *Setdb1* KO, denoted as log^2^ fold change. **I**, Heat map showing expression difference of 10 most upregulated ERVKs after *Setdb1* KO and WT ESCs, cultured in serum and 2i. Expression difference is denoted as log^2^ fold change.

To provide a comparative map of SetDB1-repressed targets in serum and 2i ESCs, we integrated our methylome analyses with publicly available H3K9me3 and H3K27me3 ChIP-seq data, both from WT and *Setdb1* KO 2i ESCs (Wu et al., 2020). In 2i ESCs, SetDB1 target regions with elevated DNAme relative to other target regions (Fig. 6D: ΔDNAme↓ SetDB1 targets) were also densely covered with H3K9me3, with H3K9me3 levels exceeding those in serum ESCs (Fig. 6E). After inducing *Setdb1* KO in 2i ESCs, these regions underwent a profound loss of H3K9me3 (Fig. 6E), despite their unaffected DNAme levels (Fig. 6D). In general, H3K27me3 levels in 2i ESCs were slightly elevated at ΔDNAme↑ SetDB1 target regions as compared to ΔDNAme↓ SetDB1 targets (Fig. 6E). However, this is unrelated to SetDB1 activity, as coverage remained similar following *Setdb1* KO. Instead, these findings likely reflect the mutual exclusivity between H3K27me3 and DNAme, the latter being more prevalent at ΔDNAme↓ SetDB1 sites.

Examining individual loci revealed that SetDB1-targeted ERVKs, primarily IAPs, were enriched with DNAme in 2i ESCs under WT conditions, albeit at much lower levels compared to serum ESCs (Fig. 6F; cf. Supplemental Fig. 3A). These findings are consistent with previous reports of partial DNAme retention at specific ERVKs in 2i ESCs (Habibi et al., 2013). Reflecting patterns observed at SetDB1 target regions, the slight DNAme enrichments at SetDB1-targeted ERVK persisted after inducing *Setdb1* KO in 2i ESCs (Fig. 6G), unlike serum ESCs (Fig. 3A). These ERVKs also exhibited higher WT H3K9me3 levels in 2i ESCs compared to serum ESCs (Fig. 6F; color scale), potentially offsetting the lack of extensive DNAme. The H3K9me3 enrichments that covered these ERVKs were erased following *Setdb1* KO. We used previously generated RNA-seq data (Wu et al., 2020) to correlate these chromatin remodellings to changes in expression levels. Upon *Setdb1* KO, SetDB1-targeted ERVKs were highly induced for expression in 2i ESCs, up to 130-fold for RLTR9B2 (Fig. 6H-I). IAP elements were also highly upregulated in *Setdb1* KO 2i ESCs, showing a 25-fold increase on average, compared to a 1.6-fold increase in *Setdb1* KO serum ESCs, consistent with previous studies (Deniz et al., 2018). Moreover, the most de-repressed ERVKs in *Setdb1* KO serum ESCs exhibited even greater de-repression in *Setdb1* KO 2i ESCs (Fig. 6I), indicating a strong reliance on SetDB1-mediated H3K9me3 silencing in the hypomethylated ground-state of pluripotency. Our findings parallel those of Sharif et al. (2016), whom showed that *Setdb1* KO in serum ESCs, combined with either *Uhrf1* (i.e. N*p95*) or *Dnmt1* KO, results in much greater derepession of repeats than *Setdb1* KO alone. Both results suggest that the absence of both DNAme and H3K9me3 yields a synergestic effect on repeat reactivation.

SetDB1-targeted genes that were demethylated in *Setdb1* KO serum ESCs (as defined in Fig. 4A) showed minimal regulation by SetDB1-mediated DNAme in 2i ESCs (Supplemental Fig. 5D). Mirroring ERVK targets, SetDB1-targeted genes were not demethylated following *Setdb1* KO in 2i ESCs. Similarly, target genes also heavily depended on SetDB1-deposited H3K9me3 in 2i ESCs, which was lost after *Setdb1* KO (Supplemental Fig. 5D) and led to strong de-repression (Supplemental Fig. 5E). Therefore, we conclude that SetDB1-dependent DNAme acts as a repressive buffer to SetDB1-mediated H3K9me3 silencing in serum ESCs, which is absent in 2i ESCs, leaving SetDB1-silenced ERVKs and genes prone to strong reactivation.

## Discussion

SetDB1 has been linked to multiple epigenetic functions in ESCs, including H3K9me3 and H3K27me3 deposition (Fei et al., 2015; Karimi et al., 2011; Matsui et al., 2010), and CTCF regulation (Warrier et al., 2022). However, the potential for direct DNAme-dependent regulation by SetDB1 and its integration with the known functions has largely remained enigmatic thus far. In this study, we addressed this gap by extensively profiling SetDB1-dependent DNAme in mouse ESCs through high-resolution methylome time course analyses following *Setdb1* KO. The methylome analyses of both serum- and 2i-cultured ESCs were integrated with ChIP-seq data of H3K9me3, H3K27me3, TET1/2, and CTCF, along with RNA-seq data, to offer a holistic view of how SetDB1-dependent DNAme intersects with the other regulatory modes. We find that ∼50% of SetDB1 targets in serum ESCs are marked by SetDB1-dependent DNAme, which is partially lost following *Setdb1* KO. These loci are also almost exclusively enriched with SetDB1-mediated H3K9me3, which is erased after KO. This tight coupling of SetDB1-mediated H3K9me3 and DNAme indicates that SetDB1 does not regulate DNAme independently, but as part of a coupled epigenetic mechanism. The H3K9me3-DNAme axis of SetDB1 is primarily used to silence ERVK-type retrotransposons, and ICRs.

Given the inseperability between SetDB1-mediated DNAme and H3K9me3 at ERVK elements, this indicates that multiple domains of SetDB1 must act in concert to establish these marks. The recruitment of SetDB1 to ERVK loci is facilitated by its triple Tudor domain (TTD), which binds to KAP1 (Zhang & He, 2024) within KRAB-ZFP-KAP1 complexes. These complexes can recognize unique DNA sequences of ERVs by incorporating distinct ZFP species (Kosugo & Hamada, 2024; Wolf et al., 2020), of which hundreds are encoded in the mouse genome (Kauzlaric et al., 2017). An interesting possibility to explain the co-regulation of H3K9me3 and DNAme is that the interaction of KRAB-ZFP-KAP1 with the TTD of SetDB1 simultaneously activates the SET domain to catalyze H3K9me3 as well as the MBD domain to recruit DNMT3 enzymes (Li et al., 2006; Markouli et al., 2021). The coupling between SetDB1-mediated H3K9me3 and DNAme could be further strengthened through DNMT1-mediated methylation, which is recruited to H3K9me3-marked sites by UHRF1 (Fang et al., 2016; Liu et al., 2013; Ren et al., 2020). This H3K9me3-DNAme silencing mechanism of SetDB1 is summarized as a model in Figure 7.

**Figure 7:**
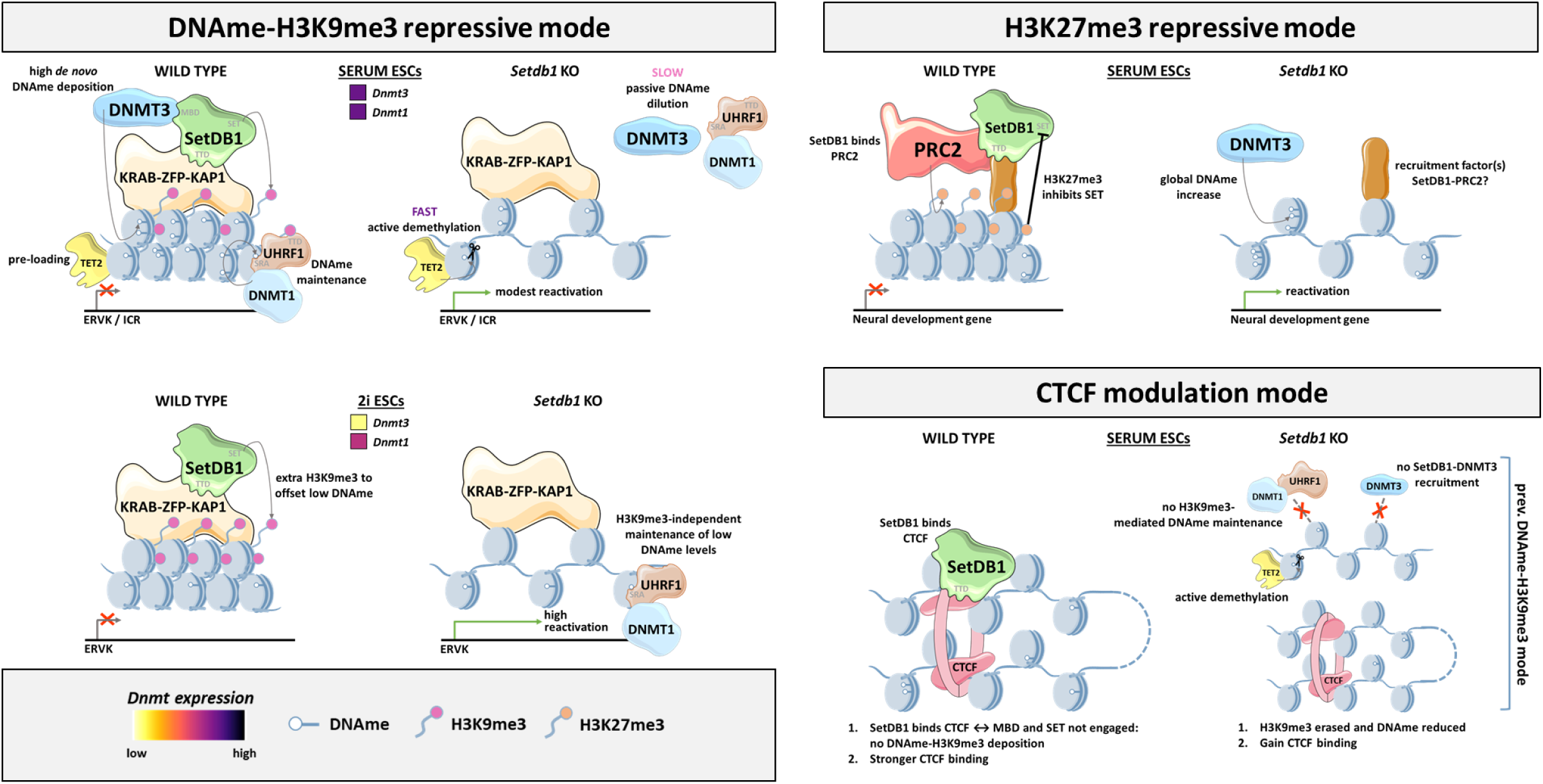
Hypothesized models of SetDB1 regulatory mechanisms in serum- and 2i-cultured mouse ESCs mediated by DNAme, H3K9me3, H3K27me3 and CTCF.

Our time-course methylome profiling provides a unique opportunity to distinguish between SetDB1 targets based on DNAme-remodeling speed. IAPs are enriched within SetDB1-targeted regions that demethylate slowly after *Setdb1* KO, while other ERVKs are found in regions that lose DNAme rapidly and extensively. This suggests that IAPs are protected from fast and extensive demethylation. Notably, we have shown that the demethylation speed in these regions is independent of their WT H3K9me3 levels and erasure after *Setdb1* KO, suggesting H3K9me3 is not the protective factor against demethylation. Instead, rapid demethylation is associated with TET pre-loading, as also previously shown for the *Dazl* locus (Leung et al., 2014). We now show that IAPs, which demethylate slowly, lack binding of TET2 enzymes in WT conditions, whereas quickly demethylated ERVKs are pre-loaded by TET2. Our finding that pre-loaded TET2 enzymes are associated with elevated 5caC levels at these loci, a marker of active and complete DNAme oxidation, supports a model in which TET2 pre-loading regulates the pace of demethylation at ERVKs. It remains to be investigated what chromatin features shield specific ERVKs from TET pre-loading and subsequent rapid enzymatic demethylation. Nevertheless, our findings suggest that SetDB1 and its recruitment of DNMT enzymes protect against demethylation in the WT situation, by counteracting the removal of DNAme by pre-loaded TET2: a safeguard lost upon *Setdb1* KO.

While SetDB1-mediated H3K9me3 and DNAme primarily mark ERVKs in serum ESCs, they also mark ICRs. Both loci are bound by SetDB1, which is recruited by KRAB-ZFP-KAP1. However, the order in which SetDB1-dependent H3K9me3 and DNAme are established at these loci appears to differ. The ZFPs that recognize ERVK and ERV1 sequences function independently of pre-existing DNAme (Rowe et al., 2013; Seah et al., 2019). We show that in WT 2i ESCs, which model pre-implantation embryos, ERVKs are densily marked with SetDB1-mediated H3K9me3 but not with SetDB1-dependent DNAme. Extensive SetDB1-dependent DNAme appears after implantation, as modelled by serum ESCs. In contrast to ERVKs, ICRs require pre-existing DNA methylation for recognition by ZFPs (e.g. ZFP57 and ZFP445), which then recruit SetDB1 to deposit H3K9me3 (Henckel et al., 2009; Liu et al., 2012; Quenneville et al., 2011; Takahashi et al., 2019). Despite differing recruitment mechanisms – i.e. methylation-sensitive for ICRs and -insensitive for ERVKs – KRAB-ZFP-KAP1 complexes guide SetDB1 to both, activating deposition of H3K9me3 and DNAme. Accordingly, we find that loss of SetDB1 in serum ESCs affects ICRs and ERVKs similarly: H3K9me3 erasure, DNAme loss, and transcriptional reactivation (Fig. 7).

Global levels of DNAme are reduced in 2i ESCs, but some SetDB1-targeted ERVKs – such as IAPs – are relatively enriched for DNAme. Unlike in serum ESCs, *Setdb1* KO in 2i ESCs does not reduce DNAme at these sites, suggesting this DNAme is maintained independently of SetDB1. We anticipate that DNMT1 preserves these intermediate levels of DNAme in 2i ESCs (Shukla et al., 2020). Potentially, even without H3K9me3 following SetDB1 loss, UHRF1 can still recognize and bind hemi-methylated DNA after replication through its SRA domain – albeit with reduced affinity – and subsequently recruit DNMT1 (Harrison et al., 2016; Liu et al., 2013; Sharif et al., 2007). UHRF1 remains nuclear and active in 2i ESCs, unlike hypomethylated PGCs (Von Meyenn et al., 2016), which would provide a means to support intermediate levels of DNAme maintenance without SetDB1-mediated H3K9me3 (Fig. 7).

Even though DNAme levels are unaffected upon *Setdb1* KO in 2i ESCs, DNAme levels remain much lower than those in *Setdb1* KO serum ESCs. Following loss of SetDB1 and SetDB1-dependent H3K9me3 in 2i ESCs, ERVKs are massively reactivated, up to 130-fold. In contrast, ERVKs are only reactivated ∼3.3-fold in *Setdb1* KO serum ESCs. This indicates that SetDB1-dependent repression is less robust in 2i ESCs, where silencing relies primarily on H3K9me3 due to the low levels of DNAme. Although reactivation of such parasitic elements potentially threatens genomic stability, the dramatic reactivation of ERVKs does not seem to account for the rapid cell death after *Setdb1* KO in 2i ESCs. Instead, loss of SetDB1 has been reported to induce expression of *Ripk3* in 2i ESCs, triggering RIPK1/3-mediated necrosome assembly and activation of the necroptosis pathway (Wu et al., 2020).

Despite its tight coupling to H3K9me3, SetDB1-mediated DNAme also coordinates the other regulatory functions of SetDB1. Fei et al. (2015) showed that SetDB1 binds to PRC2 to direct H3K27me3 deposition at a subset of binding sites where it does not catalyse H3K9me3. This aligns with our previous finding that H3K27me3 and DNAme are generally mutually exclusive (Brinkman et al., 2012; van Mierlo et al., 2019), and the observation that the H3K9me3-DNAme axis of SetDB1 and its regulation of H3K27me3 are uncoupled. Deletion of the SET domain of SetDB1 does not impair SetDB1-PRC2 interaction (Fei et al., 2015), suggesting that other domains of SetDB1 – potentially the TTD – mediate the interaction with PRC2. Thereto, PRC2 may compete with KAP1 for binding at the TTD, uncoupling the deposition of H3K27me3 and DNAme-H3K9me3. Moreover, H3K27me3 directly inhibits the SET domain of SetDB1 (Fei et al., 2015), further reinforcing this separation. Following *Setdb1* KO, SetDB1-mediated H3K27me3 is lost, leading to reactivation of proximal neural development genes (Fei et al., 2015) (Fig. 7).

SetDB1 also regulates CTCF binding – associated with active and open chromatin (Ong & Corces, 2014) – indicating functions of SetDB1 beyond heterochromatin formation. Tam et al. (2024) described that the actual loss of H3K9me3 mainly drives increased CTCF binding in *Setdb1* KO ESCs. This idea is supported by their finding that *Dnmt* triple KO ESCs, which lack DNAme but retain H3K9me3, show only minor increases in CTCF binding. Moreover, catalytic inactivation of SetDB1 (C1243A) also results in elevated CTCF binding (Tam et al., 2024). Importantly, we detect an enrichment of CTCF motifs and peaks within SetDB1 binding sites that lack DNAme, H3K9me3, and H3K27me3 enrichments. One of the more likley models based on these combined findings is that SetDB1 directly modulates CTCF when it does not engage in epigenetic silencing. This may involve TTD-mediated interactions, accounting for the uncoupling with its DNAme-H3K9me3- and H3K27me3-mediated mechanisms, which require TTD-interactions with KAP1 and PRC2, respectively (Fig. 7). Supporting this, Warrier et al. (2022) showed that deletion of SetDB1’s SET domain disrupts binding of SetDB1 to ERVs, but not to loci co-occupied by Cohesin: the topological regulator of CTCF.

Our study offers a comprehensive and dynamic characterization of SetDB1-dependent DNAme regulation during the naïve and metastable stages of pluripotency. By integrating ChIP-seq and RNA-seq datasets, we uncovered the role of SetDB1-dependent DNAme and its associated epigenetic mechanisms within the complex regulatory network of SetDB1. We cannot exclude the possibility of additional (separate or integrative) regulatory modes, such as involvement of SMAD3, whose binding motifs were highly enriched at specific SetDB1 peaks. SetDB1 may also directly regulate histone acetylation, as motifs for TFs that influence histone (de-)acetylation were identified among SetDB1 targets. Moreover, SetDB1 contains two nuclear export signal (NES) domains (Markouli et al., 2021), and recent reports suggest regulatory functions of SetDB1 outside the nucleus (Rapone et al., 2023). Further research is required to investigate these potential regulatory modes and the underlying mechanisms at the level of SetDB1’s protein domains.

## Methods

### Cell culture of mouse ESCs

*Setdb1^CKO/-^* ESCs were kindly provided by Matsui et al. (2010). ESCs were maintained in a humidified incubator at 37 °C and 5% CO_2_, and routinely tested negative for mycoplasma. Serum ESCs were cultured on 0.1% gelatin-coated culture plates in DMEM-Glutamax basal medium (Gibco) supplemented with 0.1mM non-essential amino acids (Gibco), 1mM sodium pyruvate (Gibco), 0.1mM 2-mercaptoethanol, 5·10^5^U leukemia inhibitory factor (LIF) (home-made), 10% fetal bovine serum and 25U/ml Penstrep (Gibco). 2i ESCs were maintained on 0.2% gelatin-coated culture plates in Ndiff 227 medium (Takara) supplemented with 1uM MEK inhibitor PD0325901, 3uM GSK3 inhibitor CHIR99021, together known as ‘2i’ (Ying et al., 2008), 5·10^5^U LIF (home-made) and 25U/ml Penstrep (Gibco).

### Quantitative PCR

Quantitative PCR (qPCR) was performed on extracted genomic DNA from ESCs and used as a template for amplification. Genomic DNA was isolated using phenol/chloroform extraction with ethanol precipitation. Briefly, ESCs were harvested, resuspended in SDS lysis buffer (10 mM Tris·Cl, 200 mM NaCl, 10 mM disodium EDTA, 0.5% (w/v) SDS, 100 µg/ml Proteinase K) and incubated overnight at 50 °C on a shaking heating block. The following day, an equal volume of 25:24:1 phenol:chloroform:isoamyl alcohol was added followed by vigorously inverting for 1min. Samples were centrifuged for 5min, 14,000xg, RT. The top layer was then transfered to a clean tube without disturbing the underlying organic phase. 1ml of 100% ethanol was added to the transferred top layer, after which it was vigorously inverted a few times and centrifugated for 20min, 14,000xg, 4 °C in a pre-cooled centrifuge. The supernatant was removed, and the pellet was rinsed with 1ml 70% ethanol, then centrifuged for 10min, 14,000xg, 4 °C in a pre-cooled centrifuge. The supernatant was removed and the pellet was air dried for ∼10min before dissolving. qPCR was then conducted with iQ SYBR Green Supermix (Bio-Rad) on a CFX96 Real-Time system (Bio-Rad). The DNA input was normalized to the *Act*, *Rex* and *Setdb1* exon 7 genomic loci. Primer sequences are provided in Table 1.

**Table 1:**
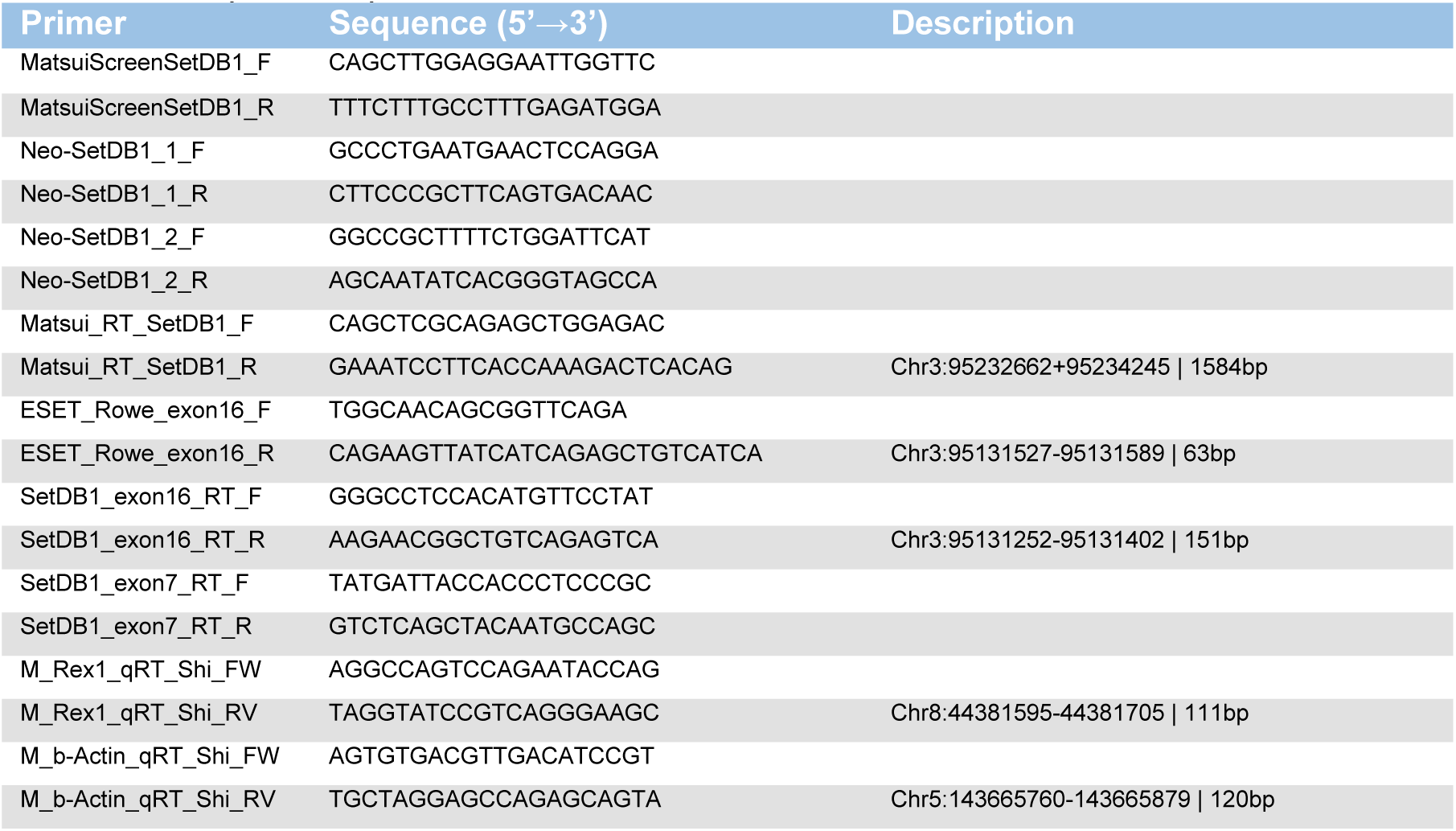
Primer sequences for qPCR.

### DNAme measurements by mass spectrometry

DNA was extracted from ESCs using the PureLink Genomic DNA Mini Kit (Invitrogen) following the manufacturer’s instructions. The samples were diluted to a final DNA concentration of 100 ng/μl, with a minimum volume of 3 μl. DNA degradation into individual nucleosides was achieved using DNA Degradase Plus (Zymo Research). These nucleosides were analyzed using high-performance liquid chromatography-tandem mass spectrometry (HPLC-MS/MS). Quantification was performed based on a standard curve generated from calibration standards. The levels of 5mC and 5hmC were expressed as percentage concentration ratios: % 5-methyl-2’-deoxycytidine to 2’-deoxyguanosine (%mdC/dG) and % 5-hydroxymethyl-2’-deoxycytidine to 2’-deoxyguanosine (%5hmdC/dG), respectively.

### WGBS & data processing

DNA from ESCs was isolated using either (i) a Cell Culture DNA Midi Kit (Qiagen), following the manufacturer’s instructions, or (ii) phenol/chloroform extraction with ethanol precipitation as described above for qPCR. WGBS on isolated DNA was conducted as previously described (Kulis et al., 2012). Libraries were sequenced on an Illumina HiSeq 2000 platform in paired-end mode (2×100 nucleotides). All WGBS (and other sequencing) analyses were performed using the *Mus musculus* NCBI GRCm39 genome assembly (mm39). Paired-end reads from the FASTQ files were trimmed using Trim Galore with the flags --paired --trim1. The trimmed reads were then aligned to the reference genome using the Bismark package in combination with Bowtie 2 (Krueger & Andrews, 2011) with the flag -n 1 --bowtie2. BAM files were deduplicated using Deduplicate Bismark with the flag -p. Cytosine methylation levels were calculated using the Bismark Methylation Extractor with the flags -p --no_overlap (Krueger & Andrews, 2011) and applying a minimum read coverage threshold of ≥5. Methylation levels of genomic features were determined by intersecting them with genomic coordinates of the mm39 reference genome, including ChIP-seq target regions, as mentioned below.

### ChIP-seq data processing

SetDB1, H3K9me3, H3K27me3, TET1, TET2, and CTCF ChIP-seq datasets were obtained from previous studies (Table 2). ChIP-seq FASTQs were processed with our in-house Seq2Science package (van der Sande et al., 2023) using the mm39 reference genome and custom ChIP-seq settings to allow read mapping against repetitive regions.

The BWA-MEM2 alignment was adjusted as follows:

|  |  |  |
| --- | --- | --- |
| 611 | remove_blacklist: false | Reads that map to blacklisted regions, i.e. highly repetitive genomic |
| 612 |  | regions, were not removed. |
| 613 | min_mapping_quality: 0 | Reads that map to repetitive regions are inherently of low quality, |
| 614 |  | hence mapping quality threshold was minimized to allow repeat read |
| 615 |  | mapping. |
| 616 | only_primary_align: false | Concerns multi-mapping reads. Typically the alignment designated |
| 617 |  | 'primary' is the best alignment, but the decision may be arbitrary, and |
| 618 |  | hence was set to 'false'. |
| 619 | remove_dups: false | Duplicate reads were kept, as these may be repetitive. |

The Macs2 settings were adjusted as follows for SetDB1 ChIP-seq peak calling:

|  |  |  |
| --- | --- | --- |
| 622 | --buffer-size 10,000 | Standard setting |
| 623 | --keep-dup 1 | Duplicate reads at the same position in paired-end data are likely |
| 624 |  | PCR artefacts, and were removed. |
| 625 | --nomodel |  |
| 626 | --extsize 100 | Normally, Macs2 builds a model based on the data itself to determine |
| 627 |  | what coordinate ranges are best applied to the peaks. However, |
| 628 |  | building a peak model on repetitive regions could be obstructed by |
| 629 |  | inherently more noise and lower mapping quality of repetitive reads. |
| 630 |  | Hence, model building was turned off, and peaks were assigned a |
| 631 |  | fixed width of 100bp centered on the peak summit. |

The Macs 2 settings were adjusted as follows for hPTM ChIP-seq peak calling:

|  |  |  |
| --- | --- | --- |
| 634 | --buffer-size 10,000 |  |
| 635 | --keep-dup all |  |
| 636 | --broad | Unlike the narrow SetDB1 ChIP-seq peaks, hPTM "peaks" likely do |
| 637 |  | not have a distinct focal point but instead resemble a broader |
| 638 |  | "blanket" pattern. Consequently, broad peak calling was applied. |
| 639 | --nomodel |  |
| 640 | --extsize 200 | The peak coordinate ranges were manually extended by an |
| 641 |  | additional 100bp beyond the SetDB1 ChIP-seq settings to resemble |
| 642 |  | a broader peak profile. |

Seq2Science’s Macs2 output provided the genomic ranges of peaks along with TMM-normalized values, which served as an approximation of peak sizes. To calculate RPKM values for each genomic feature discussed in this article, the BAM files generated by Seq2Science were imported into EaSeq (Lerdrup et al., 2016). The standard ChIP-seq import settings were used, except "only allow reads with unique positions" was disabled. In addition to intersecting the ChIP-seq data with methylome bins and SetDB1 peak regions, the data were directly mapped to the chromosomal locations of genes, ICRs (Table 3), and repeats. Repeat locations were obtained from the UCSC mm39 Repeatmask dataset. RPKM values for genomic features were calculated as follows:

- The number of overlapping DNA fragments from ChIP-targets were counted within an area ranging from the start of the regions with an offset of 0 bp, to end of the regions with an offset of 0 bp;
- Quantities were normalized to a dataset size of one million reads;
- The number of fragments was derived from the count by dividing it with (1 + DNA fragment size / size of the area).

The RPKM values were subsequently tabulated and exported from EeSeq as tab-separated TXT-files and further processed, analyzed and visualized in R (Rstudio-Team, 2022).

**Table 2:**
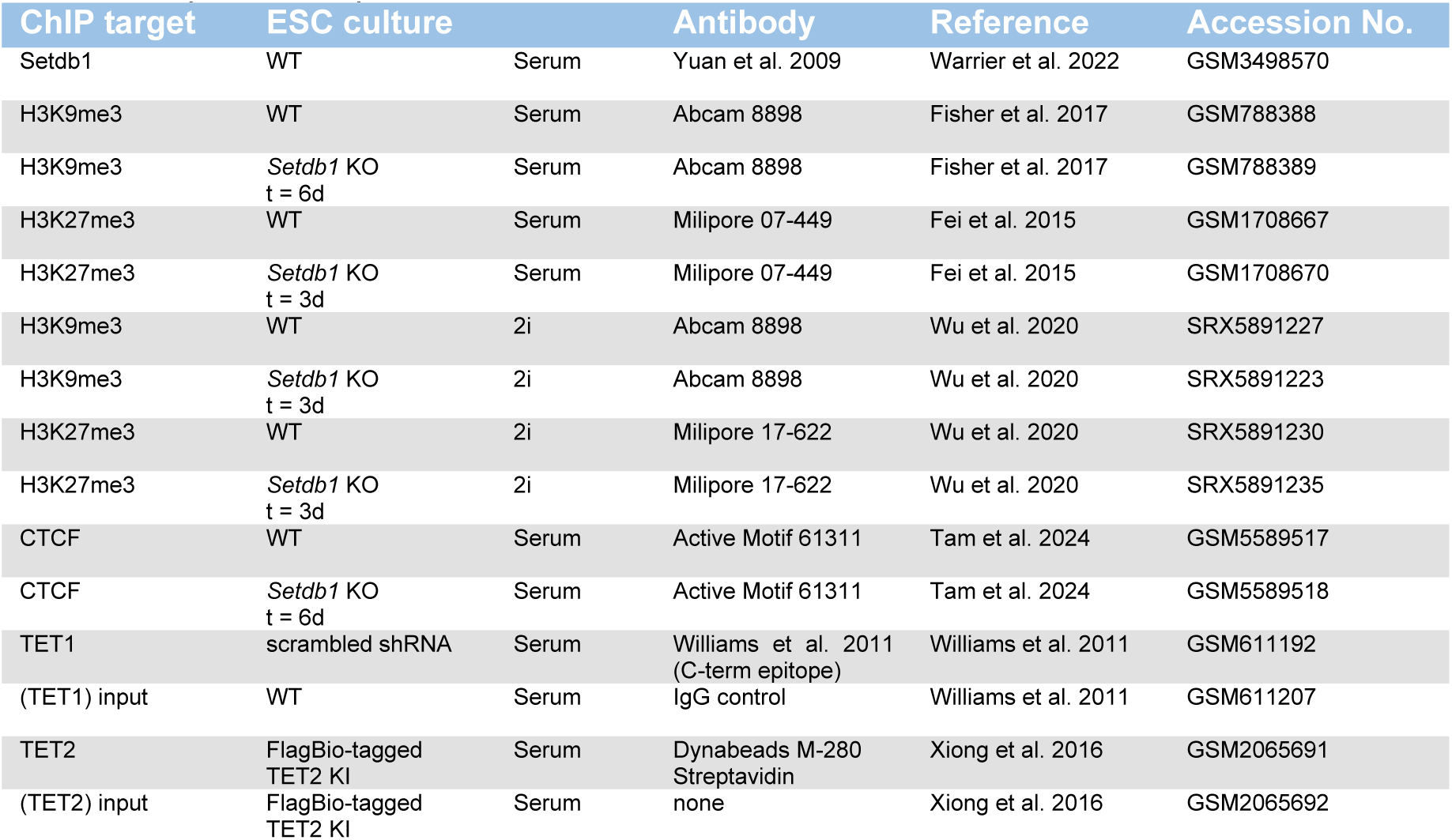
List of public ChIP-seq data used in this article.

**Table 3:**
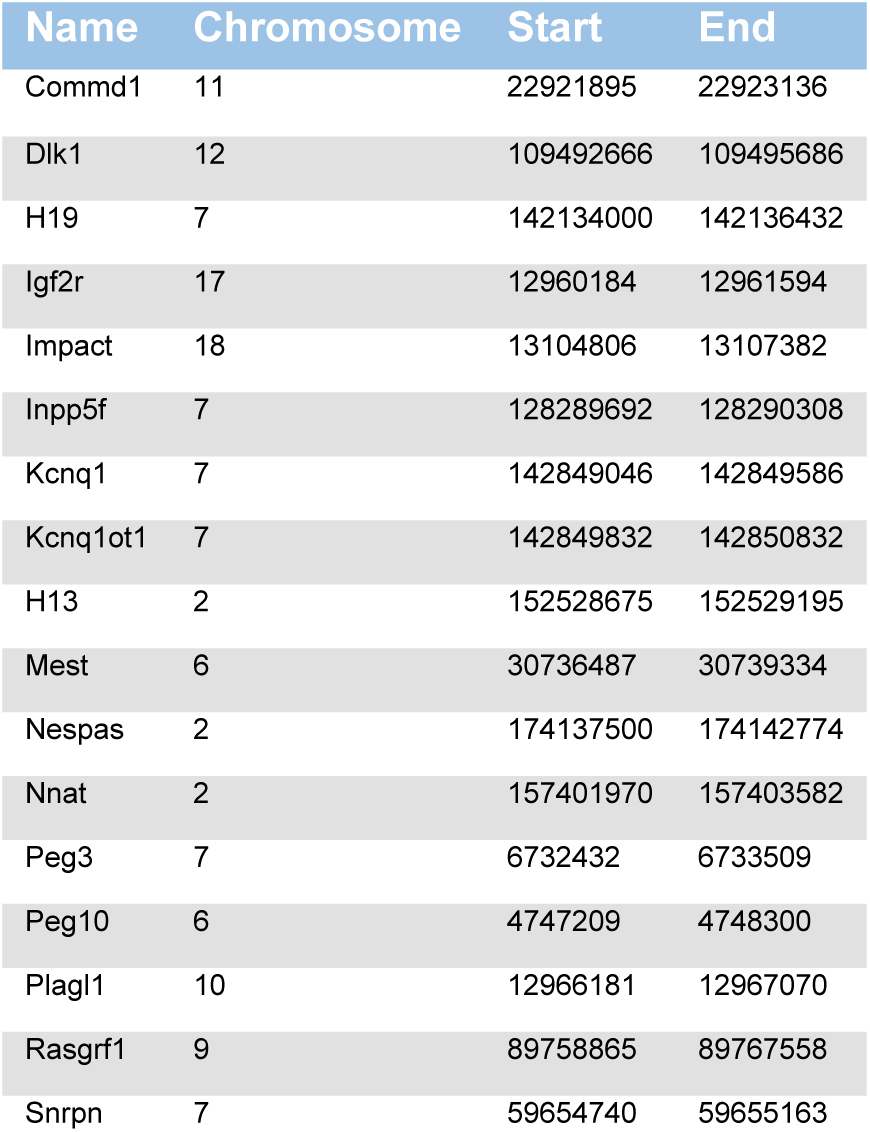
List of chromosomal locations of ICRs in mm39 reference genome.

| Name | Chromosome | Start | End |
| --- | --- | --- | --- |
| Commd1 | 11 | 22921895 | 22923136 |
| Dlk1 | 12 | 109492666 | 109495686 |
| H19 | 7 | 142134000 | 142136432 |
| Igf2r | 17 | 12960184 | 12961594 |
| Impact | 18 | 13104806 | 13107382 |
| Inpp5f | 7 | 128289692 | 128290308 |
| Kcnq1 | 7 | 142849046 | 142849586 |
| Kcnq1ot1 | 7 | 142849832 | 142850832 |
| H13 | 2 | 152528675 | 152529195 |
| Mest | 6 | 30736487 | 30739334 |
| Nespas | 2 | 174137500 | 174142774 |
| Nnat | 2 | 157401970 | 157403582 |
| Peg3 | 7 | 6732432 | 6733509 |
| Peg10 | 6 | 4747209 | 4748300 |
| Plagl1 | 10 | 12966181 | 12967070 |
| Rasgrf1 | 9 | 89758865 | 89767558 |
| Snrpn | 7 | 59654740 | 59655163 |

### Transcription factor motif analysis

Transcription factor binding motifs were identified within SetDB1 ChIP-seq peaks using Gimme Maelstrom and the non-redundant Gimme Vertebrate motif database to maximize the number of detectable motifs. Briefly, SetDB1 peaks were standardized to 200 base pairs centered on their summits, and the number of reads in the standardized peak regions were log transformed and quantile normalized. Maelstrom applies an ensemble learning approach with eight different methods combined using a rank aggregation, in order to generate motif activity z-scores, as displayed throughout the manuscript (Bruse & Heeringen, 2018). The list of transcription factor binding motifs was further curated by retaining motifs with an activity exceeding one standard deviation above the mean, while excluding duplicates, unknown transcription factors and transcription factors without mouse orthologs. Full and curated lists are available as Supplemental Table 1.

### RNA-seq data processing

RNA-seq datasets were obtained from previous studies (Table 4). Similar to ChIP-seq data, RNA-seq FASTQs were processed with our in-house Seq2Science package (van der Sande et al., 2023) using the mm39 reference genome. The reference genome pulled by Seq2Science was supplemented through -custom_annotation_extension with a BED-like GTF-file containing the chromosomal locations of all repeats in the UCSC mm39 Repeatmask dataset. Custom RNA-seq settings were applied to map reads against genes and repetitive elements, as previously reported (Teissandier et al., 2019).

The STAR alignment was adjusted as follows:

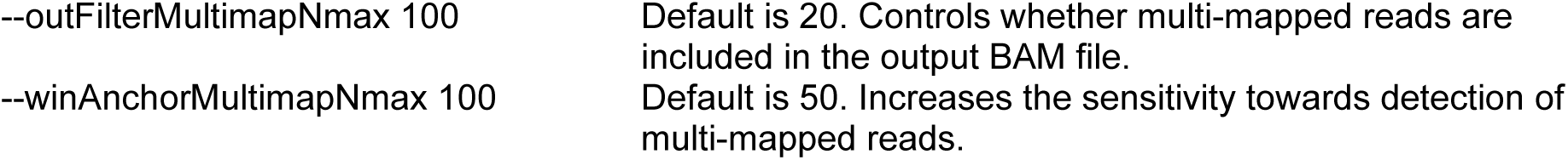

Post-alignment filtering was adjusted in the same manner as for the ChIP-seq data:

remove_blacklist: false
min_mapping_quality: 0
only_primary_align: false
remove_dups: false

The Seq2Science read count tables were filtered by omitting genes and repetitive elements with less than 3 reads. Fold-change values (WT vs *Setbd1* KO) were generated using the DESeq2 package, and further processed, analyzed and visualized in R (Rstudio-Team, 2022).

**Table 4:** List of public RNA-seq data used in this article.

| ESC culture |  | Reference | Accession No. |
| --- | --- | --- | --- |
| WT rep1 | Serum | Barral et al. 2022 | GSM5233362 |
| WT rep2 | Serum | Barral et al. 2017 | GSM5233363 |
| WT rep3 | Serum | Barral et al. 2017 | GSM5233364 |
| *Setdb1* KO rep1<br>t = 5d | Serum | Barral et al. 2017 | GSM5233365 |
| *Setdb1* KO rep2<br>t = 5d | Serum | Barral et al. 2017 | GSM5233366 |
| *Setdb1* KO rep3<br>t = 5d | Serum | Barral et al. 2017 | GSM5233367 |
| WT rep1 | 2i | Wu et al. 2020 | SRX5891234 |
| WT rep2 | 2i | Wu et al. 2020 | SRX5891239 |
| *Setdb1* KO rep1<br>t = 5d | 2i | Wu et al. 2020 | SRX5891241 |
| *Setdb1* KO rep2<br>t = 5d | 2i | Wu et al. 2020 | SRX5891248 |

## Data availability

The datasets from the whole-genome bisulfite sequencing experiments produced in this study, as FASTQ and BED files, are available at NCBI Gene Expression Omnibus **GSE299458** (https://www.ncbi.nlm.nih.gov/geo/query/acc.cgi?acc=GSE299458) with the following reviewer token: yzsxysckdradpwx.

## Acknowledgements

We thank Yoichi Shinkai for providing the *Setdb1^CKO/-^* ESCs used in this study. We also thank the RIMLS laboratory members and bioinformations for their experimental and analytical support, and Michiel Vermeulen for critically reading the draft of the paper, and communication before publication. The Marks lab is supported by a Nederlandse Organisatie voor Wetenschappelijk Onderzoek (NWO) XL grant (OCENW.XL21.XL21.100), a ZonMW Open grant (09120232310096) and a Radboud Science faculty grant (IRP voucher). The Lorincz lab is supported by a CIHR grant (PJT-190055).

## Author contributions

Conceptualization NGPB, ML, HGS, HM.

Data curation NGPB, EH.

Formal analysis NGPB, SHAH, HM.

Funding acquisition HGS, HM.

Methodology SHAH, SF, ABB, JHJ.

Project administration HM.

Supervision HGS, HM.

Visualization NGPB.

Writing – original draft NGPB, HM.

Writing – review & editing NGPB, SHAH, EH, SF, ABB, KM, JHJ, ML, HGS, HM.

---

**Supplemental Figure 1:**
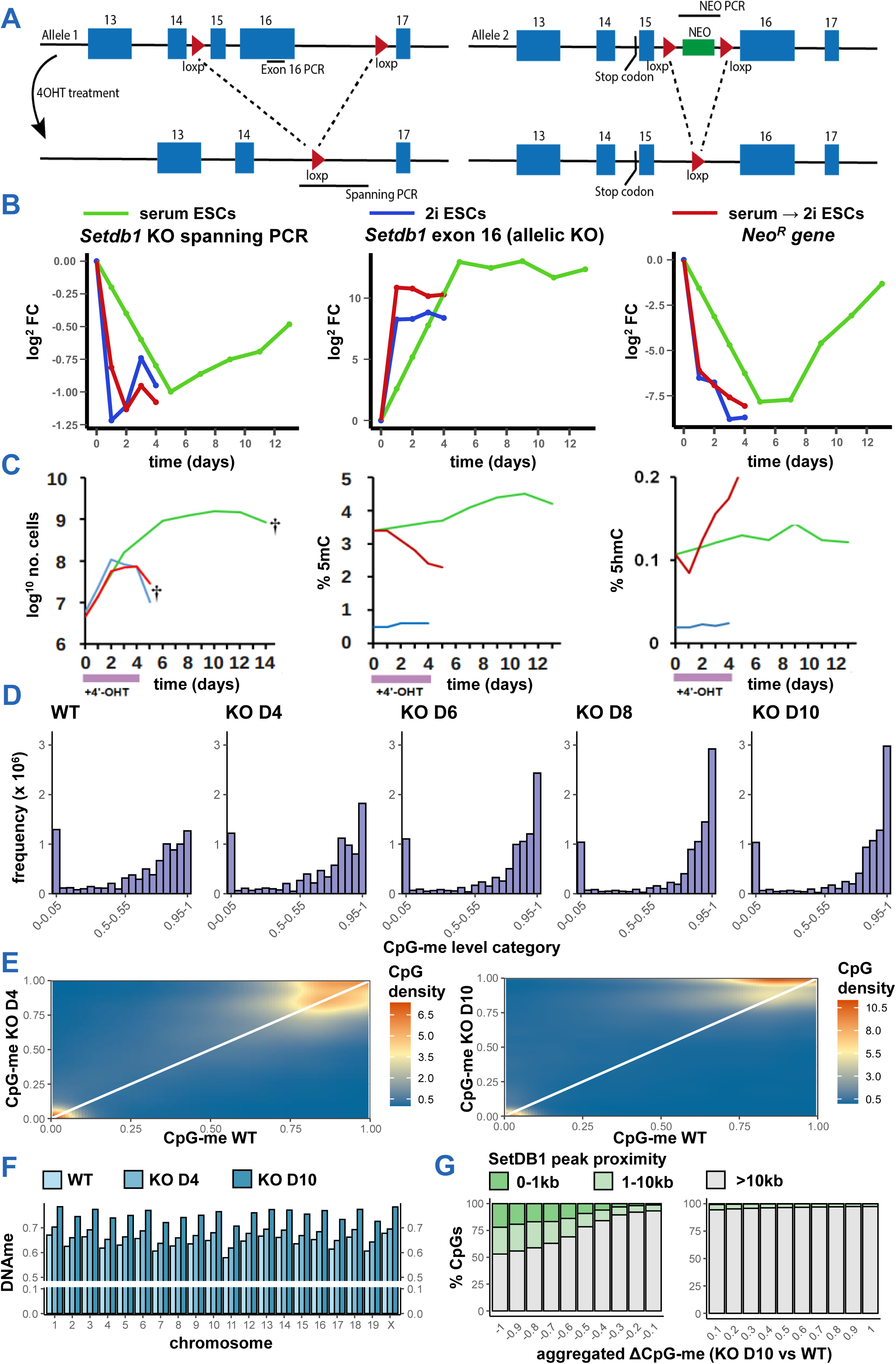
Specific SetDB1-targeted sites are demethylated following *Setdb1* KO, opposing the genome-wide DNAme increase. **A**, Schematic representation of *Setdb1* KO strategy. Left side illustrates the conditional *Setdb1* KO strategy by exon deletion through Cre-lox recombination upon 4-OHT treatment. Right side illustrates the simultaneous deletion of a neomycin resistance gene (NEO) upon 4-OHT treatment on the other allele. **B**, qPCR data confirming functionality of 4-OHT treatment to induce *Setdb1* KO showing deletion of *Setdb1* exon 15 and 16. **C**, Line graphs of cell count (left), global 5mC (middle) and global 5hmC (right) upon induction of *Setdb1* KO in serum ESCs (green line) and 2i ESCs (blue line). 5(h)mC levels were determined by mass spectrometry. Red lines represent data of *Setdb1* KO ESCs undergoing simultaneous transition from serum to 2i culture. **D**, Bar graph of CpG methylation levels in WT and *Setdb1* KO serum ESCs, as in Fig. 1B. **E**, Heatmap showing methylation remodelling of individual CpGs upon *Setdb1* KO day 4 and day 10, relative to WT serum ESCs, as in Fig. 1E. **F**, Bar graph showing average methylation of individual chromosomes in WT and *Setdb1* KO serum ESCs (WT, day 4 and day 10). **G**, Bar graph illustrating the percentage of proximally SetDB1-bound CpGs (categorized as 0-1kb, 1-10kb and >10kb distance between CpG and SetDB1 peak). CpGs were split in demethylated (left) and increasingly methylated (right) groups and subsequently ranked by aggregated ΔDNAme intervals on the horizontal axis.

**Supplemental Figure 2:**
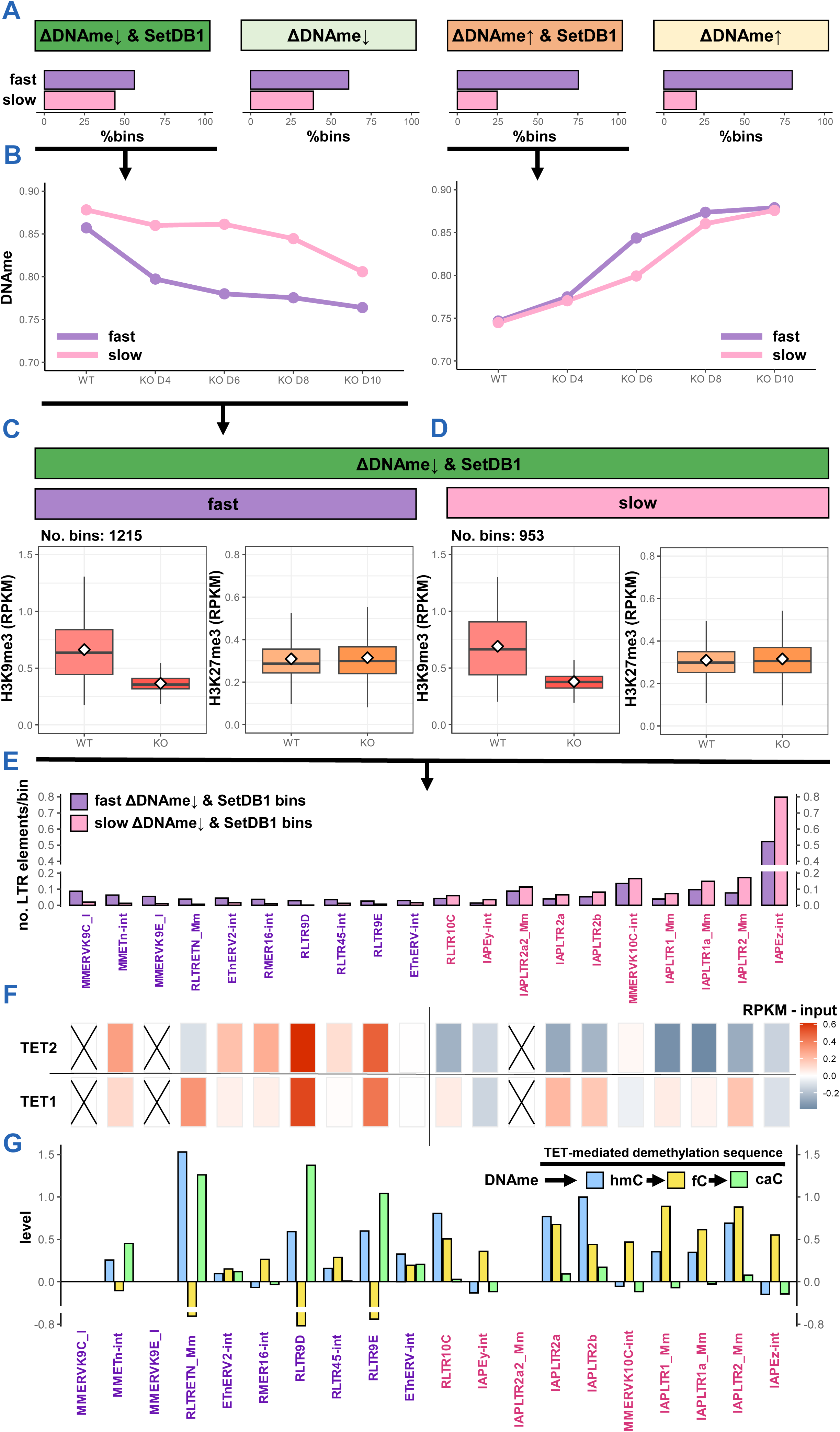
Demethylation dynamics after *Setdb1* KO of DNAme-H3K9me3-mediated targets are dependent on TET2 pre-loading. **A**, Bar graph showing the fraction of methylome bins that experience fast and slow DNAme remodelling after *Setdb1* KO in serum ESCs. **B**, Line graph showing the DNAme remodelling dynamics of fast and slow SetDB1-targeted ΔDNAme↑ and ΔDNAme↓ bins, as identified in A. Dots represent median DNAme levels over bin categories. **C-D**, Boxplots of H3K9me3 and H3K27me3 levels over entirety of bins selected from B as measured in WT and *Setdb1* KO serum ESCs. **E**, Bar graph showing the observation frequency of the 10 most-occurring ERVKs in fast and slow ΔDNAme↓ SetDB1-target bins. **F**, Heat map showing TET1 and TET2 targeting in WT serum ESCs of ERVKs listed in E. Values represent RPKM values of ChIP-seq profiles subtracted with input ChIP-seq values. Black cross indicates ChIP-seq reads could not be aligned to specific ERVK elements. **G**, Bar graph showing levels of TET-mediated oxidation products on ERVKs categorized in E-F, mined from genome-wide distribution maps of 5hmC/5fC/5caC generated with modification-specific antibodies. Levels reflect comparative measurements between TDG-deficient serum ESCs and WT ESCs.

**Supplemental Figure 3:**
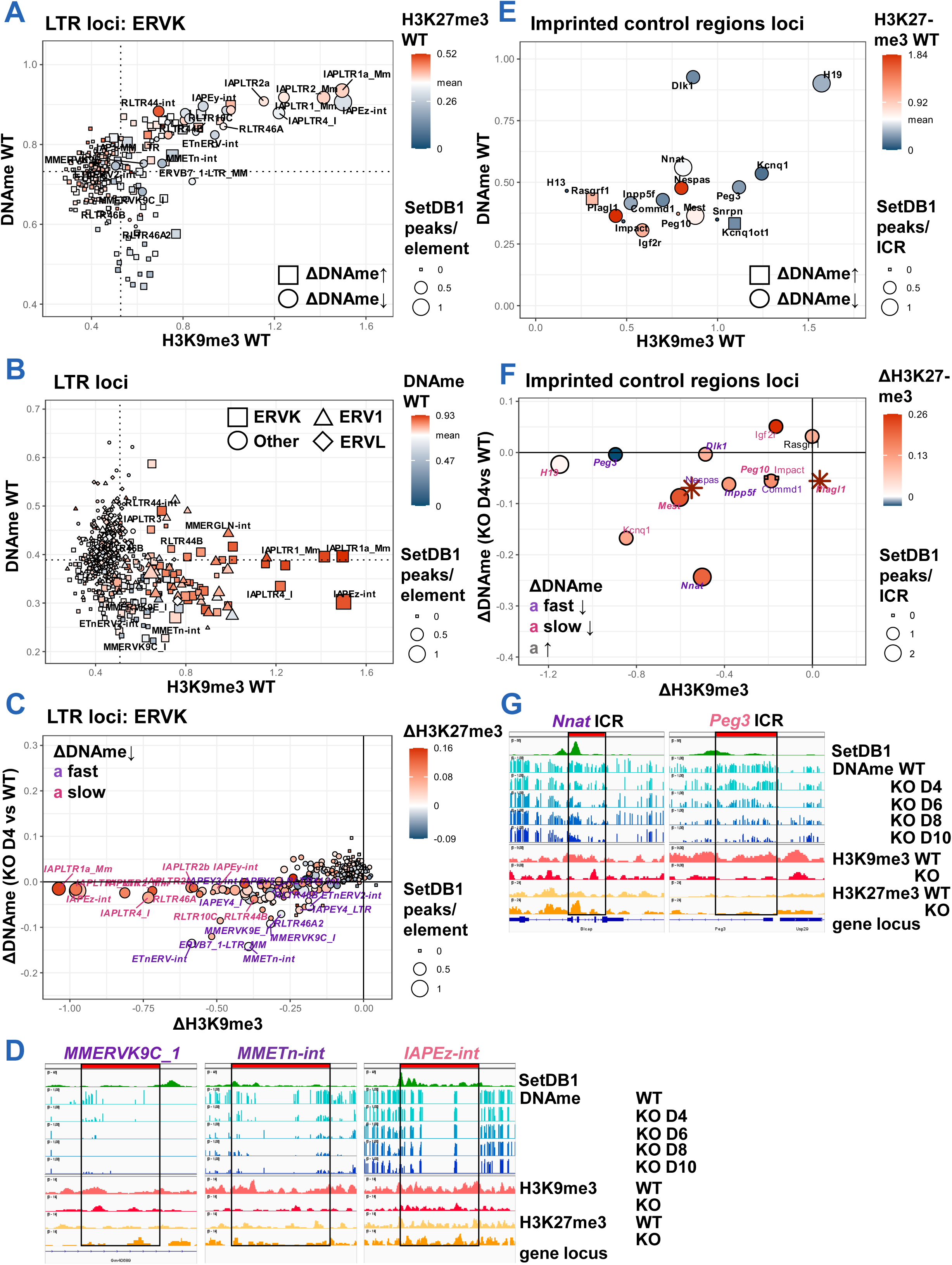
ERVK and ICR targets with SetDB1-mediated DNAme-H3K9me3 lose both repressive marks following *Setdb1* KO. **A**, Scatter plot of LTR-ERVKs showing WT H3K9me3 (ΔRPKM), DNAme (KO day 10 vs WT), and H3K27me3 (ΔRPKM) levels at ERVKs in serum ESCs. Number of SetDB1 ChIP-seq peaks per of ERVK copy (1kb proximity) displayed as point size. Dotted lines represent averages of their respectively axis parameters. **B**, Scatter plot of all LTRs, similar to A. **C**, Scatter plot similar to Fig. 3A, but vertical axis changed to ΔDNAme KO day 4 vs WT. **D**, IGV screenshots similar to Fig. 3K of MMERVK9C_1 (Chr10:70,673,637-70,675,369), MMETn-int (Chr10:83,236,248-83,240,631) and IAPEz-int (Chr11:120,212,299-120,218,751) loci. **E**, Scatter plot of ICRs, similar to A. **F** Scatter plot of ICRs, similar to C. Names in italic font if imprinted gene was transcriptionally upregulated after *Setdb1* KO. **G**, IGV screenshots of *Nnat* and *Peg3* ICR, similar to D.

**Supplemental Figure 4:**
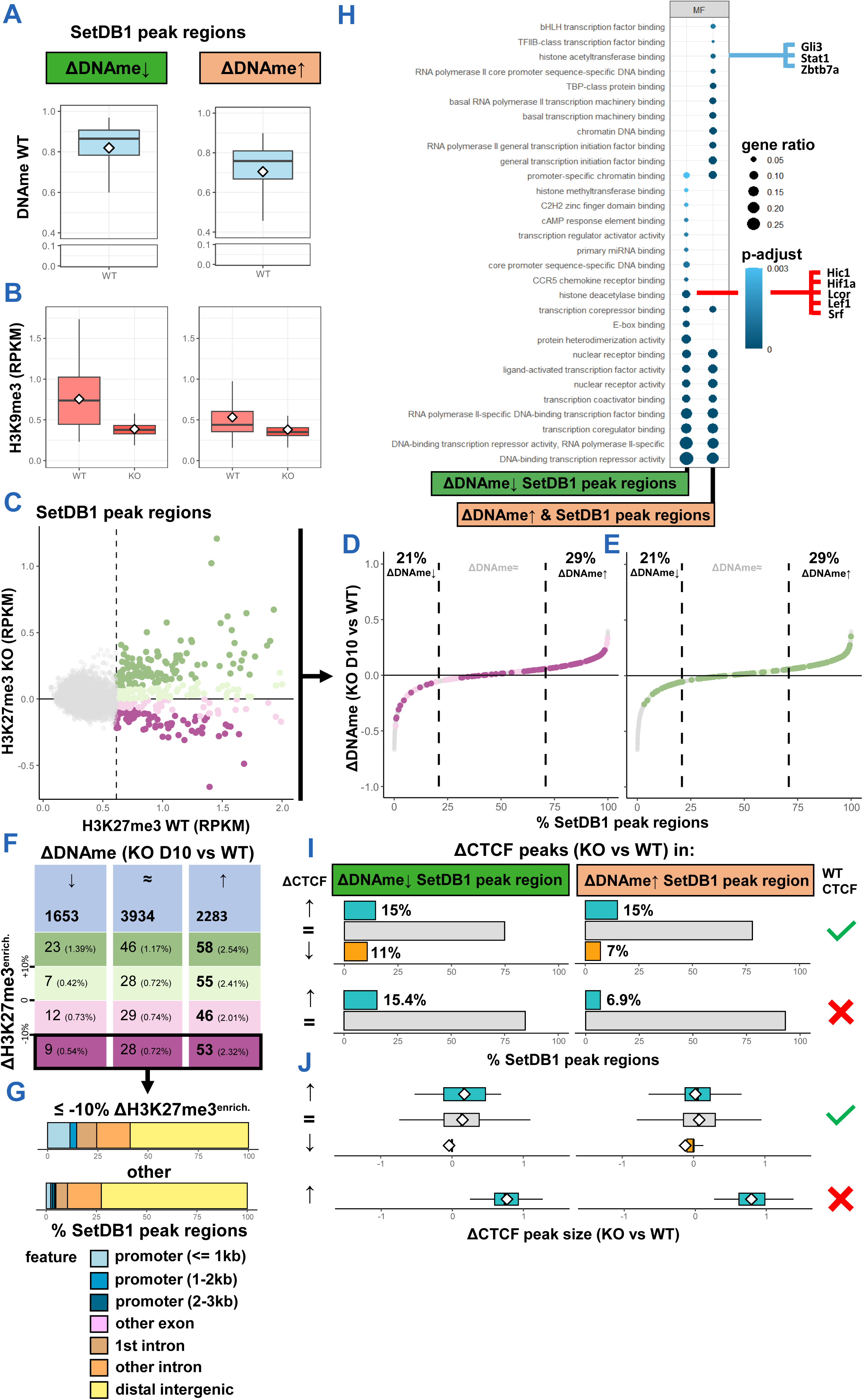
SetDB1-mediated H3K27me3- and CTCF-regulation are uncoupled from its DNAme-H3K9me3 axis. **A**, Boxplots of WT DNAme levels in serum ESCs within SetDB1 peak regions as defined in Fig. 5A. **B**, Boxplots of H3K9me3 levels in WT and *Setdb1* KO serum ESCs within SetDB1 peak regions as defined in Fig. 5A. **C**, Scatterplot of SetDB1 peak regions (10kb centered on peak summit) showing H3K27me3 in WT and *Setdb1* KO serum ESCs. Dashed vertical line represents 95th quantile. Colors match H3K27me3 remodeling category: green, > 10% H3K27me3 gain; light green, 0-10% gain; light purple, 0-10% loss; purple, >10% loss. **D-E**, Remapping of H3K27me3 remodeling categories from C to ΔDNAme-based ordering of SetDB1 peak regions of Fig. 5A. **F**, Table showing number of H3K27me3-enriched SetDB1 peak region categories from C within ΔDNAme-based categories of SetDB1 peak regions as shown in Fig. 5A. **G**, Barplot showing genomic positions of SetDB1 peaks, highlighting a subset of peaks whose regions lose H3K27me3 enrichment after *Setdb1* KO. Genomic annotations determined by ChIPseeker. **H**, Dot plot showing molecular functions of enriched TFs whose motifs are enriched in SetDB1 peak region categories from Fig. 5A, as determined by GO-analysis. **I**, Bar plots of SetDB1 peak regions, as defined in Fig. 5A and further grouped by the presence of CTCF WT peaks, showing the percentage of peak regions with changing (gain, loss) or equal (same) number of CTCF peaks upon *Setdb1* KO in serum ESCs. **J**, Boxplots showing change in average CTCF peak size between WT and *Setdb1* KO serum ESCs of categories denoted in H.

**Supplemental Figure 5:**
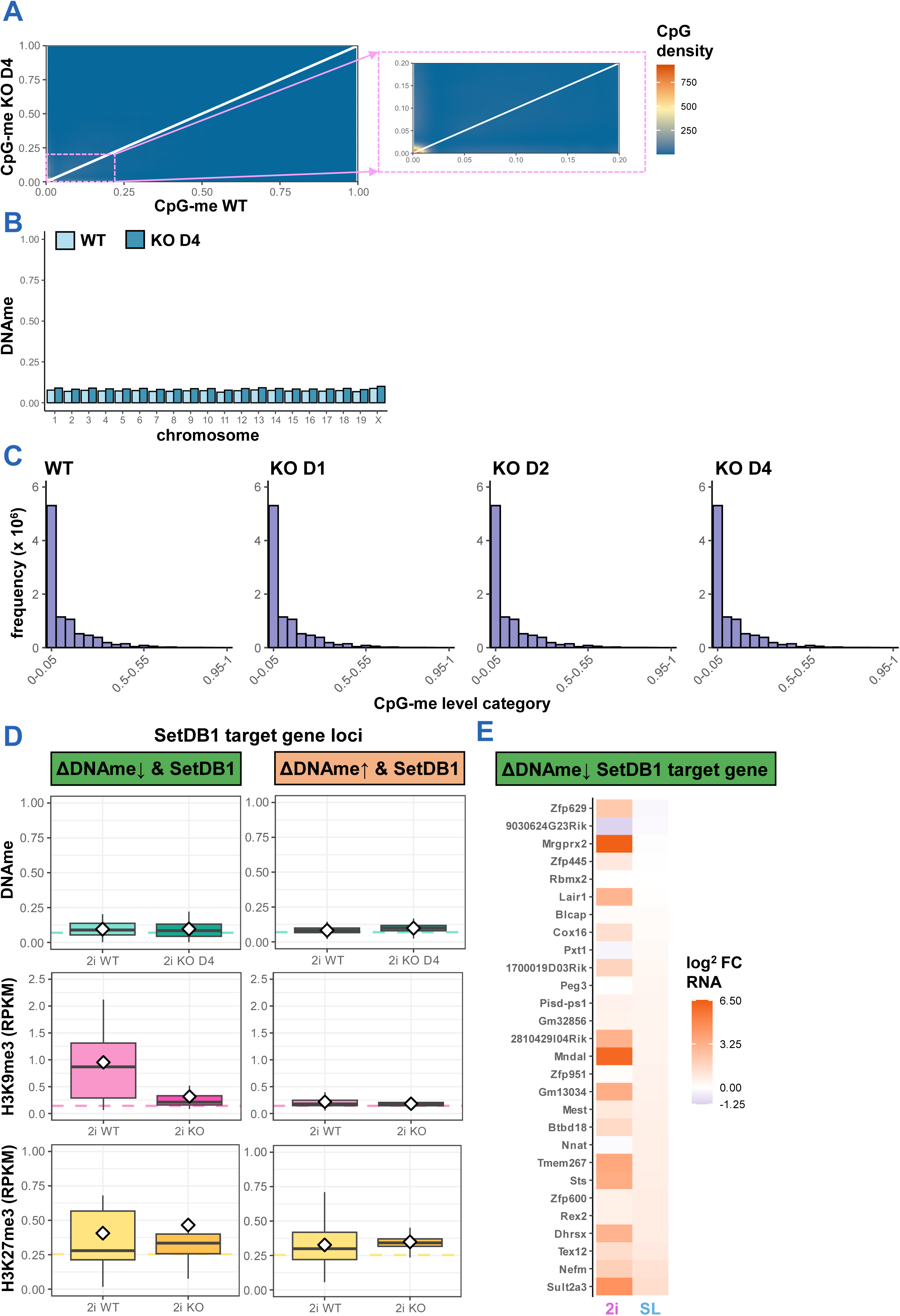
Hypomethylated 2i ESCs rely on SetDB1-mediated H3K9me3 to silence ERVK-proximal genes. **A**, Heatmaps showing methylation remodelling of individual CpGs upon *Setdb1* KO, relative to WT in 2i ESCs. CpG count density on colour scale. Vertical white lines represent slope = 1. **B**, Bar graph showing average methylation of autosomes and the X-chromosome in WT and *Setdb1* KO in 2i ESCs (WT and day 4). **C**, Bar graph of CpG methylation levels in WT and *Setdb1* KO 2i ESCs, similar to Supplemental Fig. 1D. **D**, Boxplots of DNAme, H3K9me3, and H3K27me3 levels in WT and KO 2i ESCs at SetDB1 target genes from serum ESCs (Fig. 5A: ΔDNAme↓ category). Dotted lines represent median WT levels in 2i ESCs. **E**, Heat map showing expression difference (KO vs WT) of SetDB1 target genes defined in D. Expression differences are denoted as log^2^ fold change, and shown for ESCs cultured in serum and 2i.

